# Inference of population genetic parameters from the continuously serial-sampled sequences of human seasonal influenza A/H3N2

**DOI:** 10.1101/2020.07.30.229237

**Authors:** Myriam Croze, Yuseob Kim

## Abstract

Basic summary statistics that quantify the population genetic structure of influenza virus are important for understanding and inferring the evolutionary and epidemiological processes. However, global virus sequences were sampled continuously over several decades, scattered semi-randomly over time. This temporal structure of samples and the small effective size of viral population make it difficult to use conventional methods to calculate summary statistics. Here we define statistics that overcome this problem by correcting for sampling time difference in quantifying a pairwise sequence difference. A simple method of linear regression jointly estimates the mutation rate and the level of sequence polymorphism, thus providing the estimate of the effective population size. It also leads to the definition of Wright’s *F*_ST_ for arbitrary time-series data. In addition, as an alternative to Tajima’s *D* statistic or site frequency spectrum, mismatch distribution corrected for sampling time differences can be obtained and compared between actual and simulated data. Application of these methods to seasonal influenza A/H3N2 viruses sampled between 1980 and 2017 and sequences simulated under the model of recurrent positive selection with meta-population dynamics allowed us to estimate the synonymous mutation rate and find parameter values of selection and demographic structure that fit the observation. We found that the mutation rates of HA and PB1 segments before 2007 were particularly high, and that adding recurrent positive selection in our model was essential for the genealogical structure of the HA segment. Methods developed here can be generally applied to population genetic inferences using serially sampled genetic data.

---

Of rapidly accumulating DNA sequences for evolutionary biological studies, temporal series of DNA sequences in a population are particularly valuable. While most population genetic studies indirectly infer evolutionary processes in the past from sequence diversity and polymorphism observed at present, more direct inferences are possible if sampled sequences cover a period over which the evolutionary changes of interest unfold. In particular, DNA sequences that were continuously sampled for many years from experimentally evolving organisms in laboratories (Schlötterer et al. 2014; Good et al. 2017; Van Den Bergh et al. 2018), from pathogens under surveillance by public health organizations (Holmes et al. 2016; FDA 2020), and from fossilized ancient individuals (Slatkin and Racimo 2016; www.oagr.org) can reveal their rapid evolutionary changes directly. Serially sampled sequences allow the estimation of important evolutionary genetic parameters that is not possible with data obtained at a single time point. For example, various methods were proposed to estimate the strength of natural selection and effective population size from allele frequency trajectories (Bollback et al. 2008, Steinrücken et al. 2014, Schraiber et al. 2016, Ferrer-Admetlla et al. 2016, Zinger et al. 2019). These studies assumed that nucleotide sequences are obtained from discrete time points, therefore giving discrete series of allele frequencies in a population. However, as we explain below, difficulty arises in determining allele frequencies from nucleotide sequences that were sampled continuously rather than discretely over time. Accordingly, many summary statistics that are conventionally used in population genetic inferences may not be utilized because they were primarily developed for contemporaneous genetic data (Vitti et al. 2013). This study develops evolutionary inferences that overcome such problems associated with continuously-sampled time-series genetic data, specifically as a solution to inferring complex evolutionary dynamics of seasonal influenza virus H3N2.

Being a major infectious disease agent in humans, influenza virus was heavily investigated for several decades, leading to large databases of publically available genomic sequences (GISAID and NCBI Influenza Virus Resource). Many studies demonstrated that influenza virus causing seasonal epidemics in humans, particularly subtype A/H3N2, is under strong selective pressure to evade host immune response by altering the antigenic structure of surface proteins, hemagglutinin (HA) and neuraminidase (NA) (Nelson and Holmes, 2007; Bragstad et al. 2008; Wille and Holmes, 2019). Analysing the tempo and mode of virus evolution is therefore crucial in understanding or even predicting flu epidemics. It is however challenging to correctly estimate the parameters of positive selection in influenza viruses because its complex and largely unexplained population structure, underlying seasonal epidemics over the globe, perturbs the pattern of sequence diversity from which the signatures of positive selection should be extracted. One may still use a standard approach in population genetic inference, i.e. 1) calculating various statistics for nucleotide sequence divergence and polymorphism (for example, expected heterozygosity, Tajima’s *D*, site frequency spectrum, Wright’s *F*_ST_, and coefficients of linkage disequilibrium) and 2) finding an evolutionary model that makes predictions or produces simulated data fitting those statistics, to jointly estimate the parameters of selection and population structure. However, the above statistics were originally proposed for genetic data sampled at the same time (i.e. at present) to make inference on the evolutionary past.

Global influenza viral sequences were not sampled in discrete time intervals but rather continuously over time. In addition, the effective population size of influenza virus, *Ne*, is very small, leading to a coalescent tree with short branches (two random sequences sampled in the same year coalescing within a few years back) that are connected to long “trunk” (Fitch et al. 1991, Grenfell et al. 2004; Rambaut et al. 2008). Despite such slender coalescent tree, sequence diversity is not low (mean pairwise sequence difference, π > 0.005 per nucleotide site in the HA segment) because mutation rate, μ, is large (Fitch et al. 1997; Wille and Holmes, 2019). In this case, it is not clear how one should measure the level of sequence polymorphism. As in Bedford et al. (2011) or Kim and Kim (2016), π, an estimate of 2*N*_e_μ, can be calculated for sequences sampled within a time interval of an arbitrary length. Then, the average over successive intervals can be given as a measure of sequence polymorphism. However, under the small effective population size of flu viruses, if two sequences are sampled in different time points, despite within a same interval, their expected difference can be significantly greater than 2*N*_e_μ (Figure 1A).

**Figure 1.**
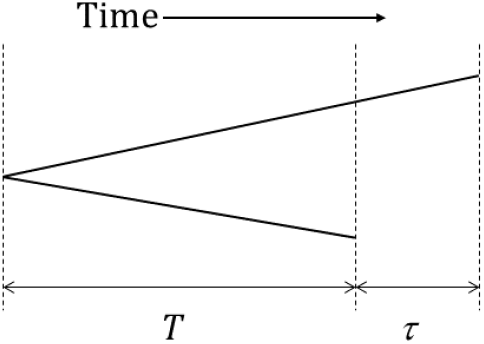
Coalescent tree of two viral sequences that are sampled at times different by τ. Assuming a constant rate μ of neutral (synonymous) mutation along a lineage, the expected neutral sequence difference is given by (2E[*T*] + τ)μ, where *T* is time to the coalescence of two contemporaneous sequences. Therefore, the expectation of synonymous sequence difference is greater than the scaled mutation rate, 2E[*T*]μ = 2*N*_e_μ, and the difference is τμ.

This problem can also be expressed as difficulty in measuring allele frequencies from serially sampled sequences. Imagine that new sequence variants either rapidly drift to extinction or increase rapidly within flu seasons, as expected for small *N*_e_, such that their frequencies are either low or high at a given time in the population. If sequences are sampled over several months during which a new variant rapidly increases from low to high frequencies, this allele is erroneously estimated to be in an intermediate frequency during this time interval, thus yielding high heterozygosity. On the other hand, if this allele started from an intermediate frequency but became fixed during the interval of sampling, allele frequency is overestimated and heterozygosity is underestimated. Therefore, to estimate allele frequency correctly, it is necessary to set time intervals for sampling as narrow as possible. However, the minimum possible length of time interval depends on how many and how evenly sequences are sampled over time per each population.

For the same reason, the conventional summaries of the pattern of sequence polymorphism, such as Tajima’s *D*, site frequency spectrum, Wright’s *F*_ST_, and the coefficients of linkage disequilibrium are difficult to obtain for continuously sampled sequences because these statistics are defined in terms of allele/haplotype frequencies at a single time point. In order to cover several decades of influenza virus evolution, one can calculate these statistics only for sequences divided into arbitrarily small time intervals, for example successive flu seasons (Bedford et al. 2011; Łuksza and Lässig, 2014). It is not clear how such time series of a statistic, which are not likely to be independent from each other, should be combined to characterize the entire genealogical structure of evolving influenza virus. A simple solution might be to obtain a single value by averaging the time series. Then again, an arbitrary decision on the number and length of sampling time intervals may greatly influence those statistics.

In this study, we show that one can systematically summarize the genealogical structure of rapidly evolving asexual population, without using arbitrary time intervals for grouping sequences, by correcting sampling-time differences in measuring nucleotide differences between sequences. In particular, we propose a time-corrected mismatch distribution (TCMD) that is an alternative to conventional statistics for quantifying genealogical structure such as Tajima’s *D* or site frequency spectrum. These approaches are used to estimate parameters of an evolutionary model of influenza virus for the HA gene sequences of H3N2 subtype sampled over decades from different geographic regions. We use the model of virus population undergoing recurrent positive selection (antigenic drift) together with complex metapopulation dynamics described in Kim and Kim (2016).

## METHODS

### Data

H3N2 viral sequences were downloaded from two databases, the Influenza Virus Resource of NCBI (Bao et al. 2008; https://www.ncbi.nlm.nih.gov/genomes/FLU/Database) and GISAID (https://www.gisaid.org). We used only genome sets, namely virus samples with all viral segments available (HA, NA, PB1, PB2, PA, NP, NS and MP). In order to avoid overlapping samples from the two databases and/or potentially extracted from the same host on the same day, we retained only one sample among those with identical base sequence, region, and date. We also discarded a sequence if it contains a notation for a base other than A, C, G and T in any position along the sequence or if the associated year, month, date, or region of extraction were not provided. We aligned only the coding region of each segments and alignment were performed with MAFFT program (Katoh et al. 2002; https://mafft.cbrc.jp/alignment/software) under default parameters. Finally, incomplete sequences for the coding region of segments were removed. At the end, we obtained a total of 6724 sequences for each segment from 1980 to 2017.

For all analyses in the following, samples from seasons 2007 to 2017 (from August 2007 to July 2017; the “10-year data”), corresponding to an alignment of 1441 sequences, were used. A season is defined as from August of a year to July of the next year. Sequences from eight regions were analysed: four from Northern Hemisphere (USA, England, China [including Hong-Kong, Macau and Taiwan] and Japan), two from Southern Hemisphere (Australia and Chile) and two from a tropical region (Nicaragua) and South East Asia (Thailand, Cambodia and Vietnam). Since the number of samples for each region and season is highly variable and uneven, we adjusted sample sizes such that our 10-year data is comparable to data sets obtained by simulations below. When there are more than 40 samples for a region in a season, we randomly subsampled 40 sequences. If there are less than 10 samples, we removed the region for the season in our analyses. At the end, we obtained 10 adjusted sets containing between 10 and 40 sequences for each region in a season. Each of these 10-year data sets yields the alignment of 999 sequences. In the following, the observed value of a test statistic or distribution is the average over these 10 data sets when it is compared to the result of simulation.

In addition, we used the alignment of 1157 sequences from 1980 to 2007 (the “27-year data”) for the joint analysis of mutation rate and nucleotide diversity. In this case, all sequences were used without dropping regions containing less than ten sequences.

### Influenza evolutionary model and computer simulation

To simulate viral sequences that are generated under the evolutionary conditions of seasonal influenza viruses, we use a metapopulation dynamic model developed by Kim and Kim (2016) in which viral sequences evolve under genetic drift, migration, and positive selection. Mimicking the HA segment of virus, a sequence has 1700 neutral (synonymous) biallelic sites, over which the level of sequence diversity is measured, and additional ε sites on which beneficial mutations increasing fitness by *s* occur. These ε sites model the “epitope” sites of HA gene that underlie the antigenic drift of influenza virus. Bidirectional mutation occurs with probability μ per generation per synonymous site and unidirectional mutation (from wild-type to beneficial) occurs with probability μ/3 per generation per epitope site. The total population is divided into eight subpopulations (demes), which undergo periodic cycles of extinction and re-colonization, connected by migration. One year is divided into 80 viral generations. Four demes model geographic regions in northern hemisphere. Their carrying capacities (*K*) reach maximum (*K*_max_) at the beginning/end of the year and remains zero between the 28^th^ and 54^th^ generation (i.e. “summer”) of a year. Two demes model those in southern hemisphere, having the opposite profile of carrying capacity reaching *K*_max_ in the middle of a year. The remaining two demes maintain constant carrying capacity *K* = 0.2*K*_max_ (the “tropical” demes). More details about migration (parameterized by *m*) and reproduction under this metapopulation dynamics are described in Kim and Kim (2016). For each set of model parameters, simulation runs for 40 years for 300 replicates and the first 30 years are removed as burn-in. Then, from the last 10 simulated years we sample around 40 sequences per deme per year.

### Basic statistics

Assume that *n* non-recombining DNA or RNA sequences were sampled over time, from a haploid population of effective size *N*_e_. As illustrated in Figure 1, synonymous difference *d_ij_* between sequences *i* and *j* with sampling time difference *τ_ij_* is expected to be E[*d_ij_*] = 2*N*_e_μ + τ_*ij*_μ, where μ is synonymous mutation rate, assuming that synonymous changes are neutral. This equation suggests that μ and *N*_e_ can be estimated by regression of *d* on τ: over the 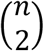 points of (τ, *d*) the slope of regression line is the estimate of mutation rate, 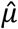. Then, samplingtime-corrected nucleotide difference between sequence *i* and *j* is given by

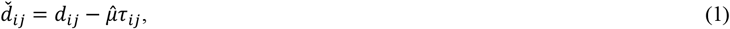

and the corrected mean pairwise sequence differences, 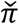, is obtained as the arithmetic mean of all 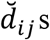 or, equivalently, as the Y-intercept of the above regression. Optionally, 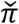 can be calculated using only the subset of sequence pairs that satisfy τ < τ_max_. Synonymous sequence differences for each pair of sequences (*d_ij_*) in a segment of H3N2 virus are obtained according to the method of Nei and Gojobori (1986) implemented in CodeML in the PAML package (Yang 1997, 2007). From 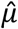 and 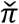, we can define the estimate of (coalescent) effective population size, 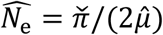.

Next, assuming that the population under consideration is subdivided into *D* demes, Wright’s *F*_ST_ can be defined for serially sampled data, by applying the above sampling-time correction to Nei’s *G*_ST_ (Nei 1973). Let *π_l_* be the arithmetic mean of 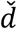 for all pairs of sequences found in deme *l*. Here, 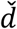 in this specific deme is still calculated using equation 1 with 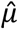 that was first determined using all sequences from all demes. Similarly, *π_lk_* is defined as the arithmetic mean of all 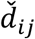 from sequence *i* in deme *l* and sequence *j* in deme *k*. Then,

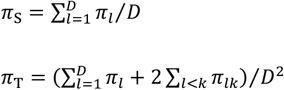

and

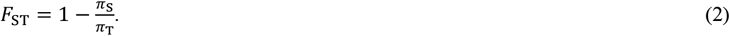

This allows one to summarize the degree of genetic differentiation between subpopulations into a single value even for data that spans a long evolutionary period. However, reminding the definition of Wright’s *F_ST_* and its genealogical interpretation (Slatkin and Hudson, 1991), we realize that πs above should quantify the level of polymorphism in sequences that are not only sampled in same subpopulations but also at the same (or nearly same) time. With limited migration, the lineages of two sequences are expected to coalesce faster if they are sampled from the same deme than from different demes. This leads to πs < π_T_ and thus *F*_ST_ > 0. If two sequences from a local population are sampled with a large time difference, however, a more recently sampled lineage is likely to have migrated to other demes by the time the other lineage is sampled. Then, their coalescent time may be no shorter than those sampled from different demes. Therefore, *F*_ST_ by equation 2 will approach to zero regardless of population subdivision as more sequence pairs with large τ are included. This is particularly true for H3N2 viruses that are globally exchanged through seasonal movement, mostly between northern and southern hemispheres. Therefore, we calculate π_S_ and π_T_ above using only sequence pairs that are sampled within τ_max_ = 300 days.

### Bootstrap tests of inter-segmental differences

Three statistics described above 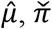, and 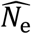, were calculated for six segments (HA, PB1, PB2, PA, NP and NA) of H3N2 data. To test if these estimates are significantly different between segments, we use bootstrap tests following the recommendation of Hall and Wilson (1991). Let 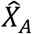 and 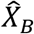 be the estimates of a parameter *X* from segment *A* and *B*. Then we define 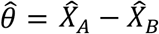 and (in case 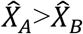 without loss of generality) test if it is significantly greater than zero. As the distribution of 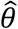 under the null hypothesis (*X_A_ = X_B_*) can be approximated by the distribution of 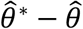, where 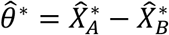 is a bootstrap value of 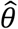, the p-value is approximately the proportion of bootstrap samples that satisfy 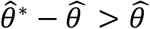. A pseudo data set is prepared by randomly sampling triplet-codon columns in the alignment of a given segment with replacement until it has the same number of codons as the original sequence. For bootstrap test, 1000 pseudo data sets for each segment pair were generated. Using the same method, we can also perform a bootstrap test to compare *X* from two different sets (e.g. old vs. recent sequences) of a single segment.

### Time-corrected mismatch distribution for HA segment

Rogers and Harpending (1992) pointed out that the distribution of pairwise sequence differences in a sample, a mismatch distribution, reflects the genealogical (coalescent) structure of sampled (non-recombining) sequences and can be therefore used to infer underlying evolutionary processes generating such a structure. We propose that the distribution of 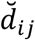 in equation 1 similarly reveals the genealogical structure of non-recombining sequences that were sampled continuously in time. We call it time-corrected mismatch distribution (TCMD). Then, TCMDs obtained from actual data and simulation are compared to find best-fit parameter values of our evolutionary model (see below). We use Kolmogorov-Smirnov test in R (Marsaglia et al. 2003) for the statistical significance of difference between TCMDs.

C codes and R scripts for the above simulation and analyses are available from https://github.com/YuseobKimLab.

## RESULTS

### Estimation of population genetic parameters

Using the method of correcting differences in sampling times described above, we jointly estimated mutation rate 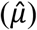 and mean pairwise synonymous difference 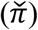 for all segments of H3N2 viruses except MP and NS that have overlapping protein-coding sequences and thus a small number of unconstrained synonymous sites. We analysed data sets of viral genomes for either 27 years (1980-2007), subsequent 10 years (2007-2017), or combined 37 years (1980-2017). In the following we simply call them 27-, 10-, or 37-year data. The estimates of mutation rates 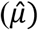 range from 1.3×10^-5^ to 3.19×10^-5^/site/day over different segments and data sets (Table 1 and Figure 2). HA segment yields the largest estimates of 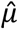 in all data sets and PB1 has the second largest for 27- and 37-year data but not for the recent 10 years. To examine if these differences in 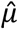 among segments are significant, we performed bootstrap tests, as described in METHODS, for all pairs of segments (Suppl. Table S1). A significant difference was not found in the recent 10 years (*p* > 0.05 for all paired comparisons). However, using 27- and 37-year samples, we find that the mutation rate of HA is significantly higher than other segments except for against PB1 segment in the data set of 37 years. In addition, we notice that 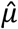 for recent 10 years is generally smaller than that for preceding 27 years. We thus applied the same bootstrap method to compare mutation rates between 10-versus 27-year data sets. We find significant difference only for HA (Figure 2).

**Figure 2.**
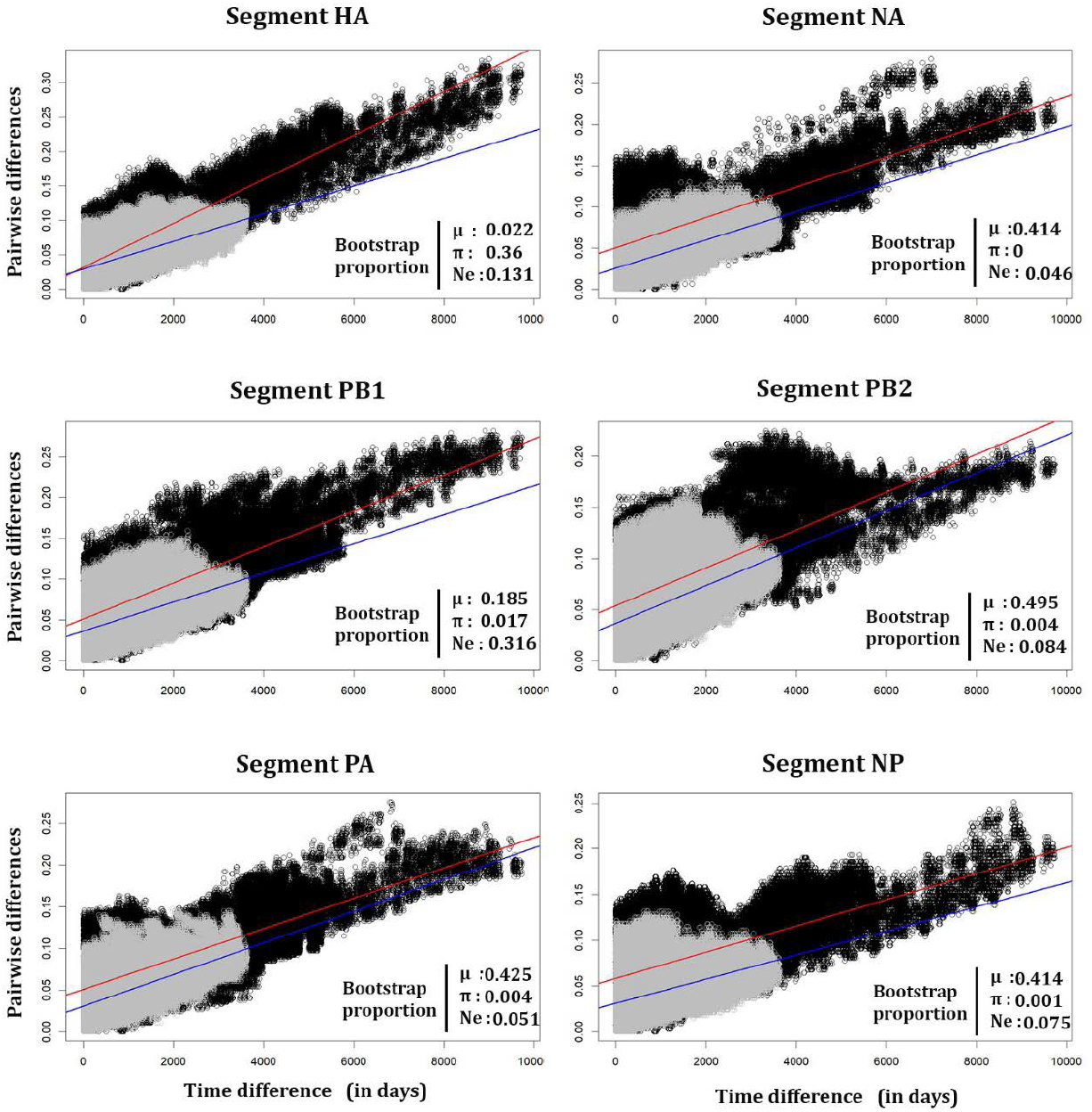
Pairwise nucleotide difference (*d*) per site of segments HA, PB1, PB2, PA, NP and NA plotted against sampling time difference (τ, in days) for H3N2 data sequences. Data points are from 27-year data (1980 to 2007; black dots) and from the 10-year data (2007 to 2017; grey dots). Regression lines for 27- and 10-year data are shown in red and blue, respectively. The proportions of bootstrap samples (in percentage) in the tests for statistical difference in 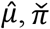, and 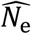 are shown below each regression plot.

**Table 1:** Sampling-time corrected, mean pairwise (synonymous) nucleotide difference, 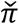, the estimate of mutation rate *μ*, and the estimate of effective population size *N*_e_ for 27-year (1980 – 2007), 10-year (2007 – 2017), and 37-year (1980 – 2017) H3N2 data sets.

| Segment | <u>27-year (1980 – 2007)</u> |  |  | <u>10-year (2007 – 2017)</u> |  |  | <u>37-year (1980-2017)</u> |  |  |
| --- | --- | --- | --- | --- | --- | --- | --- | --- | --- |
| | $\hat{\mu}$<br>($\times 10^{-5}$) | $\bar{\pi}$ | $\widehat{N_e}$ | $\hat{\mu}$<br>($\times 10^{-5}$) | $\bar{\pi}$ | $\widehat{N_e}$ | $\hat{\mu}$<br>($\times 10^{-5}$) | $\bar{\pi}$ | $\widehat{N_e}$ |
| HA | 3.19<br>(2.46/4.05)* | 0.0318<br>(0.0225/0.0427)* | 497<br>(318/730)* | 1.99<br>(1.19/2.90)* | 0.0296<br>(0.0241/0.0355) | 742<br>(502/1214) | 3.04<br>(2.38/3.77) | 0.0219<br>(0.0149/0.0288) | 361<br>(220/533) |
| PB1 | 2.20<br>(1.68/2.75) | 0.0513<br>(0.0398/0.0635) | 1165<br>(812/1674) | 1.75<br>(1.17/2.62) | 0.0366<br>(0.0315/0.0422) | 1029<br>(719/1606) | 2.50<br>(1.98/3.03) | 0.0399<br>(0.0331/0.0471) | 798<br>(619/998) |
| PB2 | 1.85<br>(1.34/2.40) | 0.0542<br>(0.0423/0.0663) | 1468<br>(1030/2152) | 1.83<br>(1.16/2.54) | 0.037<br>(0.0318/0.0419) | 1013<br>(733/1560) | 2.09<br>(1.64/2.60) | 0.0415<br>(0.0348/0.0489) | 993<br>(770/1300) |
| PA | 1.82<br>(1.30/2.37) | 0.051<br>(0.039/0.0645) | 1400<br>(923/2165) | 1.90<br>(1.19/2.68) | 0.0308<br>(0.0255/0.0366) | 812<br>(582/1270) | 1.99<br>(1.54/2.49) | 0.0396<br>(0.0320/0.0470) | 995<br>(748/1340) |
| NP | 1.43<br>(0.95/1.95) | 0.0585<br>(0.0444/0.0735) | 2050<br>(1335/3297) | 1.32<br>(0.72/2.12) | 0.0311<br>(0.0248/0.0381) | 1181<br>(763/2080) | 1.87<br>(1.38/2.41) | 0.0391<br>(0.0317/0.0470) | 1043<br>(778/1430) |
| NA | 1.84<br>(1.28/2.51) | 0.0505<br>(0.0356/0.0665) | 1372<br>(848/2087) | 1.71<br>(9.34/2.90) | 0.0256<br>(0.0207/0.0312) | 747<br>(468/1348) | 1.90<br>(1.41/2.46) | 0.0332<br>(0.0254/0.0420) | 876<br>(628/1220) |

\* Two-sided 95% confidence interval based on the distribution of estimates from 1000 bootstrap resampling.

The level of synonymous polymorphism 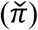 is also variable across segments. From this, we may infer the intensities of natural selection that differentially reduce the sequence diversities of segments by hitchhiking effects. However, since differences in 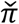 might also arise due to differences in mutation rates, we assess the effect of selection by the estimate of effective population size, 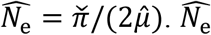 of HA segment (= 361) is smaller than the half of other segments in the 37-year data (*p* < 0.008 against NA and *p* < 0.002 against all other segments in bootstrap tests; Table 1; Table S1), as expected for this segment that is known to undergo recurrent selective sweeps (antigenic drift). Note that, as the unit of time we use (for τ and thus 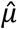) is a day, 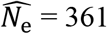 means that random two HA sequences existing at a given time during the 37 years period coalesced in one year on average. We observe similar results in the 27 years data, with 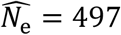. Interestingly, the effective population size of HA segment is estimated to be about twice larger for recent 10 years than estimated for the entire 37 years. 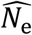 of NA is smaller than those of non-HA segments in recent 10 years but not in the entire 27 and 37 years data sets.

The above results were obtained by calculating synonymous differences for all pairs of sequences in a data set regardless of differences in sampling times (τ). Using sequence pairs that are not far apart in sampling times, i.e. τ < τ_max_ resulted in virtually identical estimates of 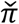 using HA sequences of the 10-year data or the corresponding simulation data (Suppl. Fig S1).

Next we calculated sampling-time corrected *F*_ST_, as describe in METHODS, for HA sequences of the 10-year data set. It slightly increases from 0.15 to 0.23 as we impose the decreasing limit of sampling time difference (τ_max_) (Suppl. Fig S1). In contrast, *F*_ST_ calculated for an equivalent 10-year data set obtained from simulation with clear population structure (Suppl. Fig. S2) increases more rapidly as we decrease τ_max_, as predicted from the nature of *F*_ST_ (see METHODS). This discrepancy between actual and simulated data occurs because H3N2 sequences are very unevenly distributed over different regions and years: as mean within-population sequence difference, π_S_, is obtained largely from pairs within a few region-year, decreasing τ_max_ does not have a large effect on π_S_. As π_S_ should be obtained in principle from pairs of sequences that existed close in time, in the following we calculate *F*_ST_ imposing τ_max_ = 300 days.

### Time-corrected mismatch distribution and population genetic inferences

As an alternative to Tajima’s *D* or site frequency spectrum that cannot be applied to a long, continuous time series of sequences, this study proposes the time-corrected mismatch distribution (TCMD; the distribution of 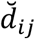) for quantifying the genealogical structure of an evolving population. We obtained TCMDs for six viral segments (two segments with insufficient number of synonymous sites, NS and MP, not included) using either 27- or 10-year data (Figure 3). In the former data set, the TCMD of the HA segment is clearly different from other segments, with more sequence pairs with small synonymous differences (Figure 3A), consistent with the significantly small estimate of effective population size (Table 1). Other segments exhibit TCMDs that are similar overall but look rather noisy, particularly towards right tails. We find, however, less fluctuation and smaller inter-segmental differences in the TCMDs of sequences from recent 10 years (Figure 3B). This is likely due to a much larger number of sampled sequences in recent years. (We averaged the TCMDs of 10 sets of subsampled sequences; see METHODS) In this recent period, NA stands out in having more sequence pairs with smaller differences. However, the difference is very small. Similar TCMDs for these six segments suggest that their evolutionary dynamics are subject to common population genetic factors, even though a certain degree of independence is expected due to reassortment between segments (Holmes et al. 2005; Berry et al. 2016). We also note in this 10-year data that the TCMD curves of multiple segments do not decrease monotonically but have small peaks or “bumps” at the right tail (close to 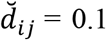). This bump is particularly prominent in the HA segment (see DISCUSSION).

**Figure 3.**
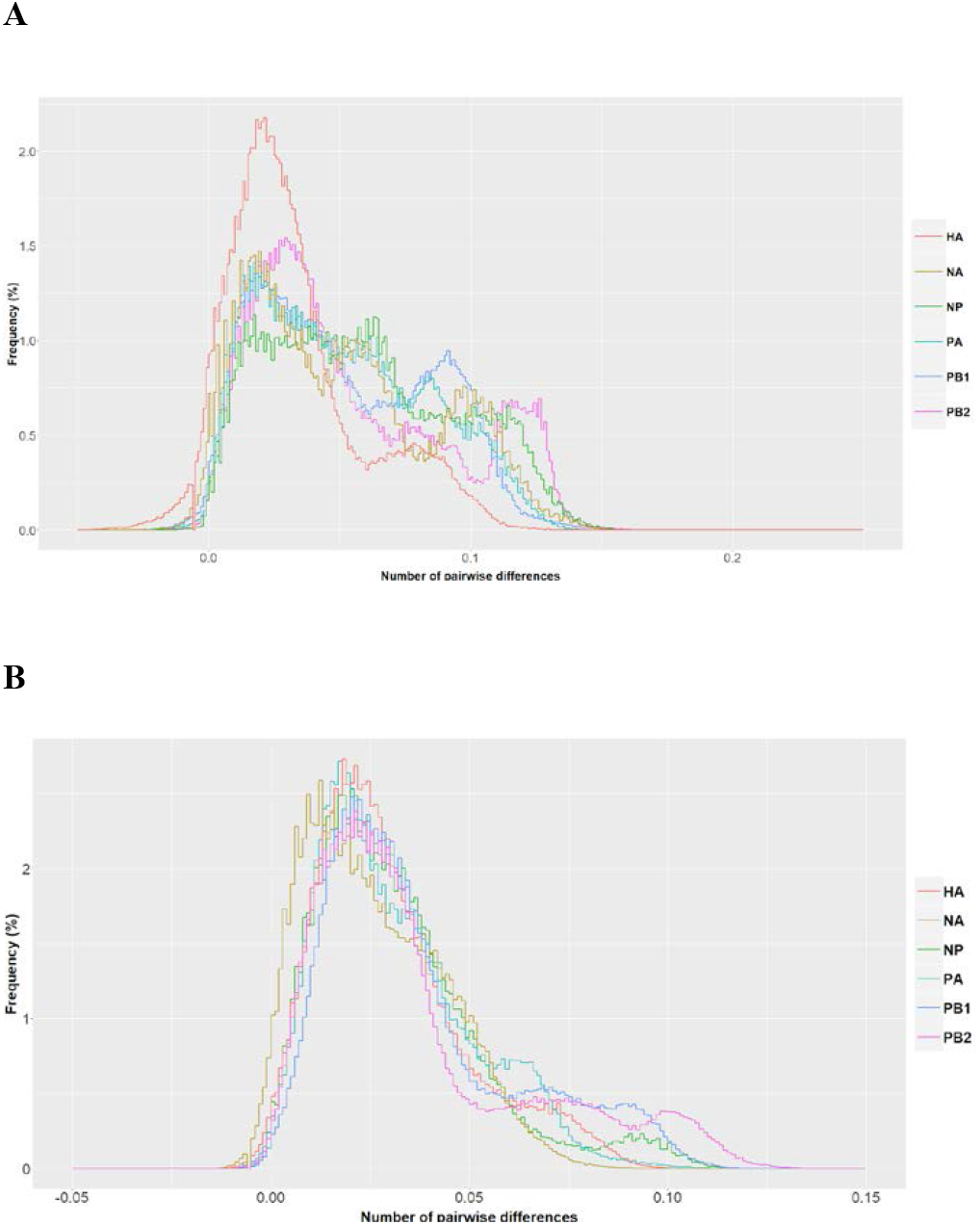
The time-corrected mismatch distributions of six influenza virus segments in the 27-year (**A**) and 10-year (**B**) H3N2 data sets. To obtain 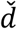 and make histograms τ_max_ = 300 and *w* = 0.002 were used.

We then ask whether summary statistics and TCMD provide sufficient information about the evolutionary processes of viruses and thus allow us to find the best-fit model and its parameter values. As a starting point, this study focuses on the model of HA sequence evolution proposed in Kim and Kim (2016) that includes the parameters of subpopulation extinction-recolonization, migration, and positive selection (see METHODS). We performed individual-based computer simulation under this model using diverse sets of parameter values to find the best match in 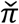, *F*_ST_ and TCMD to those from HA sequences sampled in recent 10 years.

Sequences under neutrality (ε = 0 thus no positive selection) were first generated by simulation with various combinations of *K*_max_ and *m*. However, no parameter set was found to yield 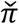 and *F*_ST_ simultaneously close to the values obtained from actual HA sequences (Suppl. Table S2). With all parameter values tried TCMD did not match that of actual data either (Figure 4). We thus tentatively conclude that demographic processes alone cannot explain the pattern of HA sequence diversity in the last 10 years.

**Figure 4.**
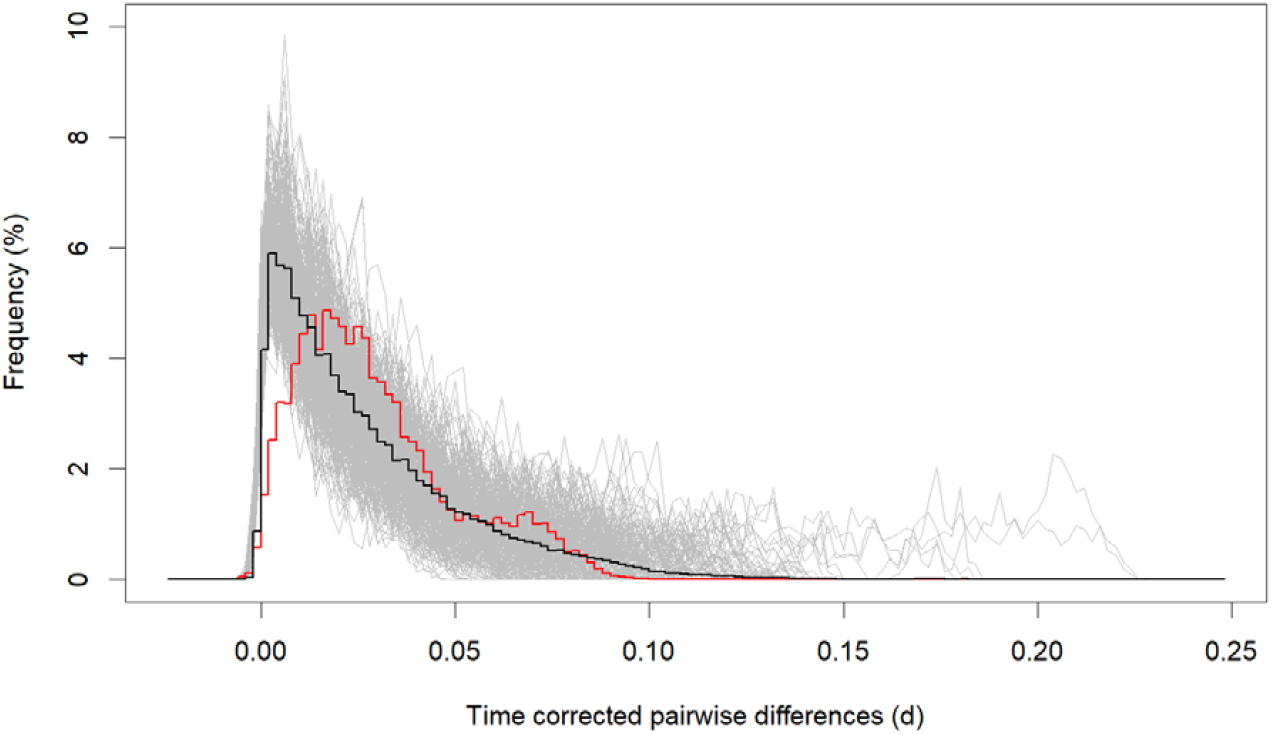
The average TCMD for simulated data under neutrality (*s* = 0) with *m* = 0.004 and *K_max_* = 110 that produce the best-fitting *F*_ST_ and 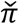 values to the observed data. To obtain 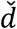 and make histograms τ_max_ = 300 and *w* = 0.002 were used.

Adding positive selection greatly improved the fit of simulation to data (Figure 5; Suppl. Fig. S3 and S4). Using *s* = 0.05 or 0.1 (values in the range that is compatible with the temporal change of allele frequencies at known antigenic variant sites; Kim and Kim 2016), we assumed the number of epitope sites ε = 10, 20, 30, or 40. Since per-site mutation rate is fixed, ε determines the rate of beneficial mutation. For each combination of *s* and ε, we first searched values of *K*_max_ and *m* until a close agreement in 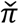 and *F*_ST_ between actual and simulated data was obtained. Then, further fine adjustments of *K*_max_ and *m* were made to find the best fit TCMD. For an increasing value of ε, the best fit was obtained with larger *K*_max_ and smaller *m* (Table 2). How responsive the shape of TCMD is to the changes of *K*_max_ and *m* is shown in Suppl. Fig. S5.

**Figure 5.**
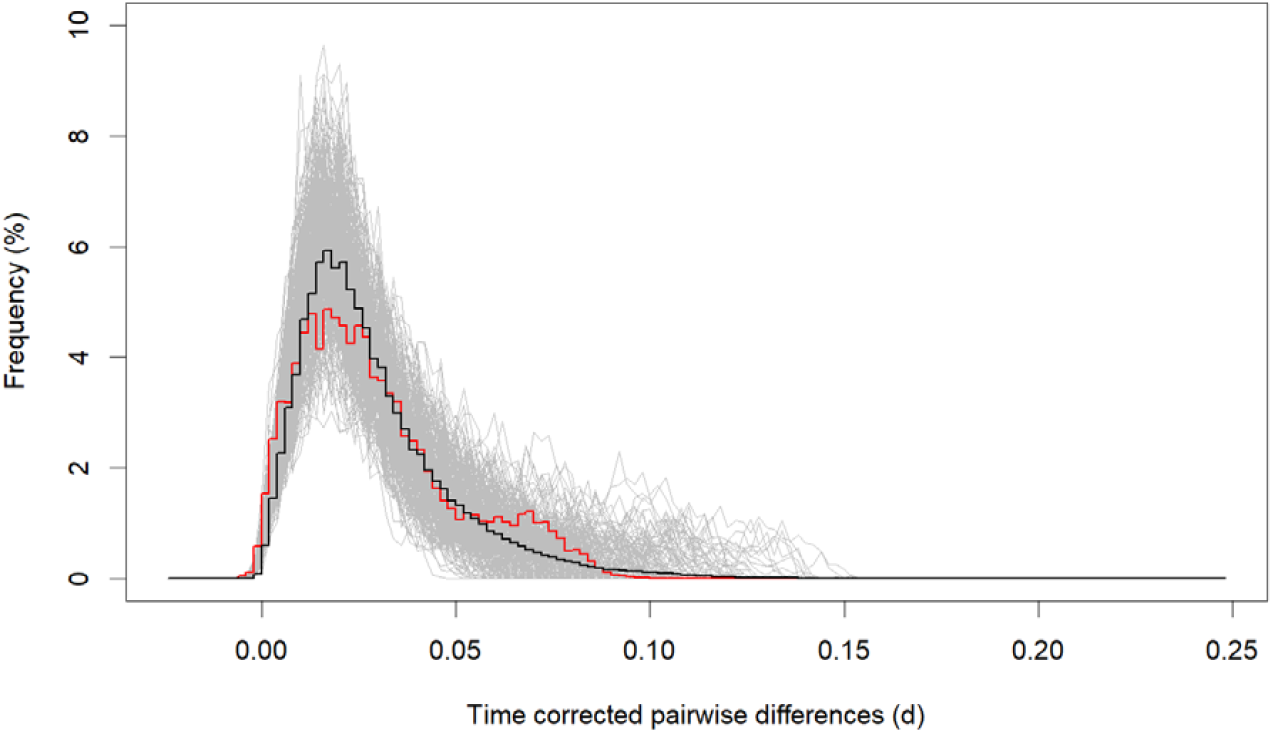
The average TCMD for simulated data under positive selection (*s* = 0.1 and ε = 10) with *m* = 0.00025 and *K_max_* = 6700, which is congruent to the TCMD of 10-year H3N2 data (Kolmogorov-Smirnov test, *p* = 0.058). To obtain 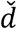 and make histograms τ_max_ = 300 and *w* = 0.002 were used.

**Table 2:**
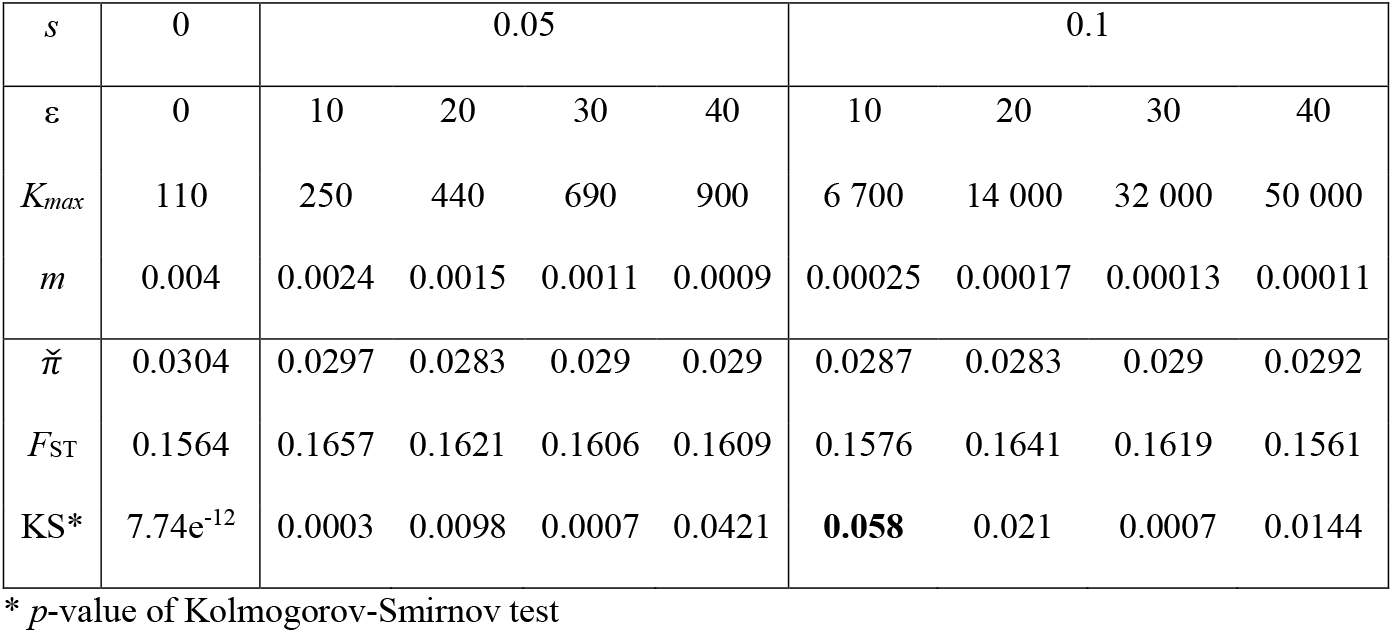
Parameters and best-fitting results of the metapopulation model under positive selection.

| $s$ | 0 | 0.05 | | | | 0.1 | | | |
| --- | --- | --- | --- | --- | --- | --- | --- | --- | --- |
| $\varepsilon$ | 0 | 10 | 20 | 30 | 40 | 10 | 20 | 30 | 40 |
| $K_{max}$ | 110 | 250 | 440 | 690 | 900 | 6 700 | 14 000 | 32 000 | 50 000 |
| $m$ | 0.004 | 0.0024 | 0.0015 | 0.0011 | 0.0009 | 0.00025 | 0.00017 | 0.00013 | 0.00011 |
| $\tilde{\pi}$ | 0.0304 | 0.0297 | 0.0283 | 0.029 | 0.029 | 0.0287 | 0.0283 | 0.029 | 0.0292 |
| $F_{ST}$ | 0.1564 | 0.1657 | 0.1621 | 0.1606 | 0.1609 | 0.1576 | 0.1641 | 0.1619 | 0.1561 |
| KS* | $7.74\text{e}^{-12}$ | 0.0003 | 0.0098 | 0.0007 | 0.0421 | **0.058** | 0.021 | 0.0007 | 0.0144 |

\* $p$ -value of Kolmogorov-Smirnov test

The quantitative measurement of agreement between the TCMDs of simulation (averaged over replicates) and data was given by the *p-value* of Kolmogorov-Smirnov (KS) test, the null hypothesis of which assumes the identity of two distributions. We found *p* > 0.05 (thus agreement between data and simulation) for one parameter set: *s* = 0.1, ε = 10, *K*_max_ = 6700 and *m* = 0.00025 (Figure 5). The second best fit (*p* = 0.041) was obtained with *s* = 0.05, ε = 40, *K*_max_ = 900 and *m* = 0.0009). The best fit TCMDs for each set of *s* and ε (Table 2) are shown in Figures S3 and S4. For a given *s*, the best-fit TCMDs appear to be similar across different values of ε upon visual inspection, necessitating the quantitative evaluation of fits by the KS test. To obtain TCMDs here, for both simulated and actual data, we used 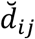 for pairs whose sampling time differences are less than τ_max_ = 300 days and the bin size *w* = 0.002. Changes from this scheme uniformly increase or decrease *p*-values in the KS tests. However, we consistently found the above sets of parameter values the best-fitting ones.

Further simulations were performed with less frequent beneficial mutations (ε = 2 and 5) and adjusting other parameters to generate best-fitting 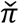 and *F*_ST_. As expected, TCMDs obtained from these simulations are intermediate between that of best-fit neutral model and those of selection with frequent positive selection (ε ≥ 10) (Suppl. Fig. S6). Therefore, TCMD was highly informative in detecting the presence of positive selection but of rather limited use in inferring ε. Namely, with ε varying in the range of large values (> 10), the genealogical structure a segment in our model, captured by TCMD, could be made to fit the HA sequence data reasonably well by adjusting other parameters.

We also tried simulations with *s* = 0.15 but a close fit was obtained only with very large values of *K*_max_ (> 50,000). Since a previous study found that the effective population size of H3N2 (under which new beneficial alleles arise) is small enough to suppress soft selective sweeps (Kim and Kim 2016) we consider such large values of *K*_max_ unrealistic. Moreover it is very time consuming (taking several weeks for one parameter set) to simulate for so large *K*_max_. Consequently, we decided not to continue analysis for *s* = 0.15.

## DISUSSION

In this study, we used continuously serial-sampled sequences of influenza A/H3N2 viruses from public databases (GISAID and NCBI) to calculate the mutation rates for different segments (HA, NA, PA, NP, PB1 and PB2). We calculated synonymous differences (*d_ij_*), with sampling time difference τ_*ij*_, in all pairs of sequences in our data sets and used the linear regression to estimate the synonymous mutation rate. The method of linear regression has been used before to calculate viral substitution rate (e.g. in HIV-1 Lukashov and Goudsmit, 1998), but all sequences were compared to the common ancestor or oldest sequence. However, such an approach is prone to error as it ignores the coalescent tree linking the oldest and other sequences in the sample (Figure 1). By explicitly incorporating this coalescent process in our linear regression we could not only estimate mutation rate but also jointly estimate the mean pairwise sequence differences and thus the effective population size.

We estimated mutation rates to range from 1.3×10^-5^ to 3.19×10^-5^/site/day. These results are similar to the estimates of synonymous substitutions rate found by previous studies, for example Hanada et al. (2004) which estimated the average synonymous substitution rate of 6.84×10^-3^/site/year (≈ 1.87×10^-5^/site/day) when all segments of Human Influenza A virus were combined. Interestingly we found that the estimate of synonymous mutation rate on HA between 1980 and 2007 is significantly higher, relative both to the estimates on other segments in the same period and to the estimate on the same segment between 2007 and 2017. Significant difference between the two periods is not observed in other segments. PB1 shows a similar trend in 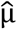, although the estimate in the 27-year data is significantly higher only against NP (Table 1). While it was clearly demonstrated that the HA gene evolves rapidly under recurrent positive selection on nonsynonymous variants (i.e. antigenic drift; Nelson and Holmes, 2007; Rambaut et al. 2008; Bhatt et al. 2011), the elevation of synonymous substitution rates in this gene needs a further explanation. One possibility is that synonymous sites on the influenza viral genome are not completely neutral but under weak constraints resulting from codon usage bias or the formation of secondary structure in the single-stranded RNA segment, which slows down the substitution rate relative to the rate expected under complete neutrality. Then, recurrent positive selection on linked nonsynonymous sites interferes with such weak selection (i.e. reduces effective population size) and can therefore increase the rate of substitutions synonymous sites (Sharp and Li 1987). This hypothesis implies that recurrent positive selection on HA was more frequent between 1980 and 2007 than between 2007 and 2017. Indeed, significantly smaller estimate of effective population size for the HA segment in 1980-2007 than in 2007-2017 (Table 1) is compatible with more frequent positive selection in the former period.

In addition, it is noted that the HA and PB1 segments came from avian influenza virus and all other segments from human H2N2 at the origin of H3N2 virus (Wendel et al. 2015; Allen and Ross, 2018). Then, evolutionary change at HA and PB1 might have been faster during the earlier period as they had to adapt to a new host, which can explain lower effective population sizes and higher synonymous mutation rates on these segments before than after 2007. Alternatively, codon usages, secondary structures, or G+C content of HA and PB1 might have evolved faster, directly increasing the rates of synonymous substitutions, in the earlier period in their adaptation to new host (dos Reis et al. 2009). Indeed, we find that the G+C content of HA decreased over time and the rate of change was greater before than after 2007 (Figure S7). We also utilized sampling-time corrections for obtaining an *F*_ST_ statistic and mismatch distributions to measure the genealogical structure of a population without being affected by how sampled sequences are scattered over time. Our *F*_ST_ statistic is moderately dependent on τ_max_, which should be set minimum in principle. In practice, if multiple sequences from each of demes under consideration are available at similar time points, say less than three months apart for seasonal influenza viruses, this restriction on τ_max_ should not be a problem. Our estimate of *F*_ST_ ≈ 0.16 confirms the earlier results that, despite rapid global circulation, HA sequences of H3N2 viruses are genetically differentiated over geographic regions. It indicates that despite rapid viral transmission across regions (“migration” rate in our model) genetic drift within each region is relatively stronger.

Modifying the mismatch distribution of Rogers and Harpending, we proposed the TCMD as an alternative to Tajima’s *D* statistic or site frequency spectrum that were defined for sequences sampled at one time point (or at nearly identical time points in the coalescent time scale). A TCMD captures the genealogical structure of population and thus provides information for finding the fitting parameters of a population genetic model. Although TCMD itself is not a single summary statistic, congruence or distance between TCMDs, particularly between those of actual and simulated data, can be easily measured for parameter estimation. However, as our individual-based simulation for the joint demographic-selection model of H3N2 virus is highly time-consuming, the entire parameter space of the model could not be systematically explored, for example through approximate Bayesian computation. We found the best-fit TCMD for each set of chosen *s* and ε by making fine adjustments of demographic parameters (*K*_max_ and *m*) only. Therefore, further simulations will probably reveal different sets of parameters that yield better agreement to the observed data. Even further improvement in inference will likely depend on proposing a qualitatively different model that incorporates important evolutionary processes determining the evolutionary dynamics of H3N2 viruses not considered here. In fact, we find from the visual inspection of evolutionary trees for HA sequences in the last 10 years that a considerable proportion of sequence pairs are highly divergent (thus the “bump” near the right tail of TCMD; Figure 3). This might indicate the presence of an evolutionary force to promote coexistence of two divergent HA lineages.

Methods developed here can be generally applied to serially-sampled population genetic data. We inferred population genetic parameters separately for each segment of influenza virus, which, not undergoing recombination, is a natural unit of independent evolution. However, our methods including TCMD must be applicable to recombining sequences as well. We think that correct inference can be made if simulation can generate data with the matching rate of recombination. This can for example lead to the evolutionary analysis of other RNA viruses undergoing homologous recombination, such as SARS-CoV-2 whose sequences were also continuously serial-sampled.

## ACKNOWLEDGEMENT

This research was supported by the National Research Foundation (NRF) grants 2018M3A9H4055197 and 2015R1A4A1041997 funded by the Korean government.

## Supplementary data

**Table S1.** Bootstrap tests for the significance of difference in mutation rates, pairwise sequence diversity, and effective population size between segments, for 27-, 10-, and 37-year data sets. Each number is the percentage of resampled data that yield the difference of a given statistic twice larger than the difference in the observed data.

|  |  | 27-year (1980-2007) |  |  | 10-year (2007-2017) |  |  | 37 year (1980-2017) |  |  |
| --- | --- | --- | --- | --- | --- | --- | --- | --- | --- | --- |
| | | $\hat{\mu}$ | $\tilde{\pi}$ | $\widehat{N_e}$ | $\hat{\mu}$ | $\tilde{\pi}$ | $\widehat{N_e}$ | $\hat{\mu}$ | $\tilde{\pi}$ | $\widehat{N_e}$ |
| HA vs | PB1 | **0.026** | **0.005** | **0.007** | 0.326 | **0.040** | 0.138 | 0.099 | **0** | **0.002** |
|  | PB2 | **0.012** | **0** | **0.007** | 0.379 | **0.037** | 0.165 | **0.019** | **0** | **0.002** |
|  | PA | **0.004** | **0.012** | **0.014** | 0.425 | 0.359 | 0.359 | **0.006** | **0.001** | **0.001** |
|  | NP | **0** | **0.002** | **0.018** | 0.010 | 0.359 | 0.134 | **0.007** | **0** | **0.020** |
|  | NA | **0.008** | **0.025** | **0.011** | 0.294 | 0.153 | 0.478 | **0.007** | **0.020** | **0.080** |
| PB1 vs | PB2 | 0.175 | 0.347 | 0.204 | 0.438 | 0.482 | 0.430 | 0.149 | 0.390 | 0.133 |
|  | PA | 0.152 | 0.484 | 0.280 | 0.391 | 0.084 | 0.208 | 0.090 | 0.457 | 0.153 |
|  | NP | **0.017** | 0.225 | 0.068 | 0.178 | 0.106 | 0.356 | 0.059 | 0.406 | 0.140 |
|  | NA | 0.168 | 0.488 | 0.281 | 0.476 | **0.004** | 0.154 | 0.057 | 0.115 | 0.362 |
| PB2 vs | PA | 0.467 | 0.345 | 0.415 | 0.445 | 0.057 | 0.239 | 0.382 | 0.378 | 0.473 |
|  | NP | 0.114 | 0.329 | 0.155 | 0.138 | 0.092 | 0.314 | 0.294 | 0.324 | 0.442 |
|  | NA | 0.481 | 0.353 | 0.408 | 0.411 | **0.001** | 0.184 | 0.316 | 0.077 | 0.292 |
| PA vs | NP | 0.133 | 0.190 | 0.128 | 0.119 | 0.482 | 0.169 | 0.390 | 0.433 | 0.446 |
|  | NA | 0.479 | 0.510 | 0.488 | 0.358 | 0.093 | 0.398 | 0.425 | 0.134 | 0.286 |
| NP vs | NA | 0.167 | 0.241 | 0.132 | 0.241 | 0.086 | 0.154 | 0.469 | 0.158 | 0.247 |

**Table S2.** Sampling-time-corrected sequence diversity and *F*_ST_ in the simulation of metapopulation dynamics under neutrality (*s* = 0). The values in bold are the closest to the observation. τ_max_ = 300 days.

| $K_{\max}$ | | $m$ | | | | |
| --- | --- | --- | --- | --- | --- | --- |
|  |  | 0 | 0.00001 | 0.0001 | 0.001 | 0.01 |
| 100 | $\tilde{\pi}$ | NA | 0.237 | 0.141 | 0.025 | 0.064 |
| | $F_{ST}$ | NA | 0.952 | 0.916 | 0.459 | 0.0291 |
| 250 | $\tilde{\pi}$ | 0.255 | 0.243 | 0.128 | **0.043** | 0.064 |
| | $F_{ST}$ | 0.895 | 0.938 | 0.828 | **0.320** | 0.029 |
| 500 | $\tilde{\pi}$ | 0.259 | 0.248 | 0.100 | 0.064 | 0.112 |
| | $F_{ST}$ | 0.803 | 0.924 | 0.675 | 0.225 | 0.014 |
| 1,000 | $\tilde{\pi}$ | 0.268 | 0.255 | 0.107 | 0.104 | 0.179 |
| | $F_{ST}$ | 0.647 | 0.887 | 0.542 | 0.132 | 0.005 |
| 2,000 | $\tilde{\pi}$ | 0.284 | 0.254 | 0.139 | 0.154 | 0.267 |
| | $F_{ST}$ | 0.387 | 0.818 | 0.423 | 0.074 | 0.005 |
| 5,000 | $\tilde{\pi}$ | 0.320 | 0.262 | 0.222 | 0.262 | 0.368 |
| | $F_{ST}$ | -0.105 | 0.643 | 0.272 | 0.033 | 0.003 |
| 10,000 | $\tilde{\pi}$ | 0.362 | 0.282 | 0.292 | 0.338 | 0.425 |
| | $F_{ST}$ | -0.544 | 0.502 | 0.176 | 0.015 | 0.000 |

**Table S3.**
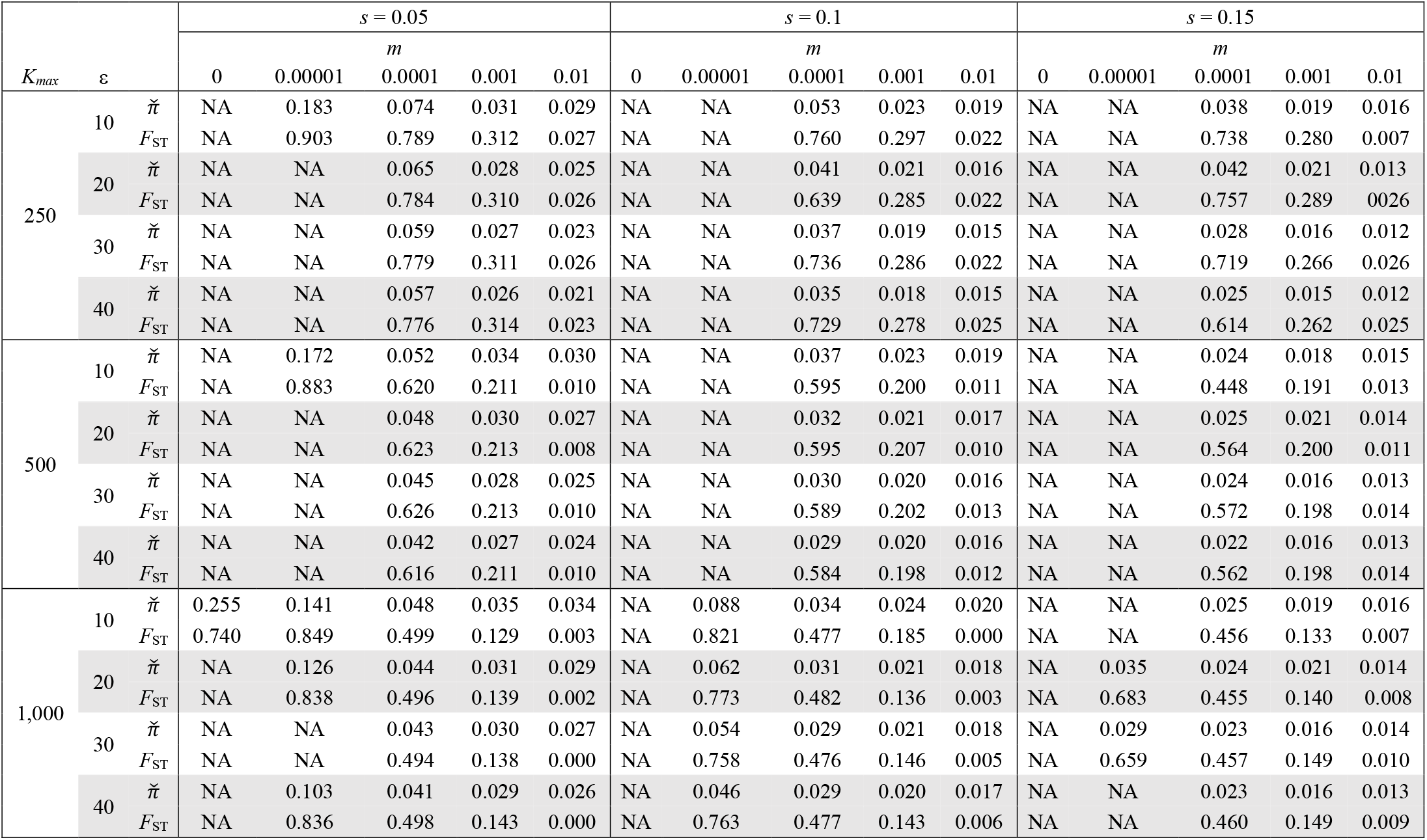
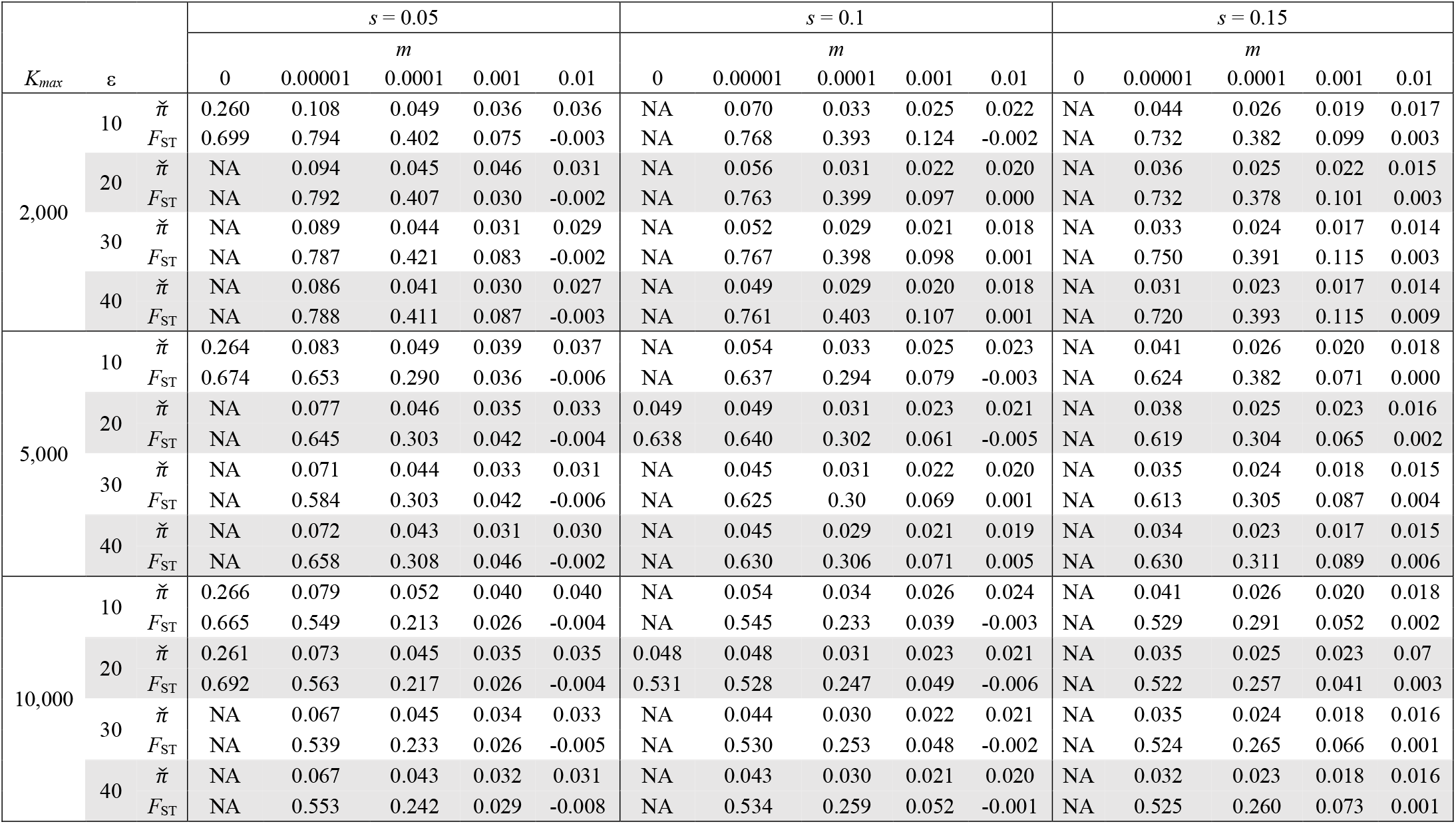
Sampling-time-corrected sequence diversity and *F*_ST_ in the simulation of metapopulation dynamics with positive selection (τ_max_ = 300 days)

| $K_{\max}$ | $\epsilon$ | | $s = 0.05$ | | | | | $s = 0.1$ | | | | | $s = 0.15$ | | | | |
| --- | --- | --- | --- | --- | --- | --- | --- | --- | --- | --- | --- | --- | --- | --- | --- | --- | --- |
| | | | 0 | 0.00001 | $m$<br>0.0001 | 0.001 | 0.01 | 0 | 0.00001 | $m$<br>0.0001 | 0.001 | 0.01 | 0 | 0.00001 | $m$<br>0.0001 | 0.001 | 0.01 |
| 250 | 10 | $\tilde{\pi}$ | NA | 0.183 | 0.074 | 0.031 | 0.029 | NA | NA | 0.053 | 0.023 | 0.019 | NA | NA | 0.038 | 0.019 | 0.016 |
| | | $F_{ST}$ | NA | 0.903 | 0.789 | 0.312 | 0.027 | NA | NA | 0.760 | 0.297 | 0.022 | NA | NA | 0.738 | 0.280 | 0.007 |
| | 20 | $\tilde{\pi}$ | NA | NA | 0.065 | 0.028 | 0.025 | NA | NA | 0.041 | 0.021 | 0.016 | NA | NA | 0.042 | 0.021 | 0.013 |
| | | $F_{ST}$ | NA | NA | 0.784 | 0.310 | 0.026 | NA | NA | 0.639 | 0.285 | 0.022 | NA | NA | 0.757 | 0.289 | 0.026 |
| | 30 | $\tilde{\pi}$ | NA | NA | 0.059 | 0.027 | 0.023 | NA | NA | 0.037 | 0.019 | 0.015 | NA | NA | 0.028 | 0.016 | 0.012 |
| | | $F_{ST}$ | NA | NA | 0.779 | 0.311 | 0.026 | NA | NA | 0.736 | 0.286 | 0.022 | NA | NA | 0.719 | 0.266 | 0.026 |
| | 40 | $\tilde{\pi}$ | NA | NA | 0.057 | 0.026 | 0.021 | NA | NA | 0.035 | 0.018 | 0.015 | NA | NA | 0.025 | 0.015 | 0.012 |
| | | $F_{ST}$ | NA | NA | 0.776 | 0.314 | 0.023 | NA | NA | 0.729 | 0.278 | 0.025 | NA | NA | 0.614 | 0.262 | 0.025 |
| 500 | 10 | $\tilde{\pi}$ | NA | 0.172 | 0.052 | 0.034 | 0.030 | NA | NA | 0.037 | 0.023 | 0.019 | NA | NA | 0.024 | 0.018 | 0.015 |
| | | $F_{ST}$ | NA | 0.883 | 0.620 | 0.211 | 0.010 | NA | NA | 0.595 | 0.200 | 0.011 | NA | NA | 0.448 | 0.191 | 0.013 |
| | 20 | $\tilde{\pi}$ | NA | NA | 0.048 | 0.030 | 0.027 | NA | NA | 0.032 | 0.021 | 0.017 | NA | NA | 0.025 | 0.021 | 0.014 |
| | | $F_{ST}$ | NA | NA | 0.623 | 0.213 | 0.008 | NA | NA | 0.595 | 0.207 | 0.010 | NA | NA | 0.564 | 0.200 | 0.011 |
| | 30 | $\tilde{\pi}$ | NA | NA | 0.045 | 0.028 | 0.025 | NA | NA | 0.030 | 0.020 | 0.016 | NA | NA | 0.024 | 0.016 | 0.013 |
| | | $F_{ST}$ | NA | NA | 0.626 | 0.213 | 0.010 | NA | NA | 0.589 | 0.202 | 0.013 | NA | NA | 0.572 | 0.198 | 0.014 |
| | 40 | $\tilde{\pi}$ | NA | NA | 0.042 | 0.027 | 0.024 | NA | NA | 0.029 | 0.020 | 0.016 | NA | NA | 0.022 | 0.016 | 0.013 |
| | | $F_{ST}$ | NA | NA | 0.616 | 0.211 | 0.010 | NA | NA | 0.584 | 0.198 | 0.012 | NA | NA | 0.562 | 0.198 | 0.014 |
| 1,000 | 10 | $\tilde{\pi}$ | 0.255 | 0.141 | 0.048 | 0.035 | 0.034 | NA | 0.088 | 0.034 | 0.024 | 0.020 | NA | NA | 0.025 | 0.019 | 0.016 |
| | | $F_{ST}$ | 0.740 | 0.849 | 0.499 | 0.129 | 0.003 | NA | 0.821 | 0.477 | 0.185 | 0.000 | NA | NA | 0.456 | 0.133 | 0.007 |
| | 20 | $\tilde{\pi}$ | NA | 0.126 | 0.044 | 0.031 | 0.029 | NA | 0.062 | 0.031 | 0.021 | 0.018 | NA | 0.035 | 0.024 | 0.021 | 0.014 |
| | | $F_{ST}$ | NA | 0.838 | 0.496 | 0.139 | 0.002 | NA | 0.773 | 0.482 | 0.136 | 0.003 | NA | 0.683 | 0.455 | 0.140 | 0.008 |
| | 30 | $\tilde{\pi}$ | NA | NA | 0.043 | 0.030 | 0.027 | NA | 0.054 | 0.029 | 0.021 | 0.018 | NA | 0.029 | 0.023 | 0.016 | 0.014 |
| | | $F_{ST}$ | NA | NA | 0.494 | 0.138 | 0.000 | NA | 0.758 | 0.476 | 0.146 | 0.005 | NA | 0.659 | 0.457 | 0.149 | 0.010 |
| | 40 | $\tilde{\pi}$ | NA | 0.103 | 0.041 | 0.029 | 0.026 | NA | 0.046 | 0.029 | 0.020 | 0.017 | NA | NA | 0.023 | 0.016 | 0.013 |
| | | $F_{ST}$ | NA | 0.836 | 0.498 | 0.143 | 0.000 | NA | 0.763 | 0.477 | 0.143 | 0.006 | NA | NA | 0.460 | 0.149 | 0.009 |

**Table S3 (next).** Sampling-time-corrected sequence diversity and $F_{ST}$ in the simulation of metapopulation dynamics with positive selection ( $\tau_{\max} = 300$ days)
| | | | $s = 0.05$ | | | | | $s = 0.1$ | | | | | $s = 0.15$ | | | | |
| --- | --- | --- | --- | --- | --- | --- | --- | --- | --- | --- | --- | --- | --- | --- | --- | --- | --- |
| | | | $m$ | | | | | $m$ | | | | | $m$ | | | | |
| $K_{\max}$ | $\varepsilon$ | | 0 | 0.00001 | 0.0001 | 0.001 | 0.01 | 0 | 0.00001 | 0.0001 | 0.001 | 0.01 | 0 | 0.00001 | 0.0001 | 0.001 | 0.01 |
| 2,000 | 10 | $\tilde{\pi}$ | 0.260 | 0.108 | 0.049 | 0.036 | 0.036 | NA | 0.070 | 0.033 | 0.025 | 0.022 | NA | 0.044 | 0.026 | 0.019 | 0.017 |
| | | $F_{ST}$ | 0.699 | 0.794 | 0.402 | 0.075 | -0.003 | NA | 0.768 | 0.393 | 0.124 | -0.002 | NA | 0.732 | 0.382 | 0.099 | 0.003 |
| | 20 | $\tilde{\pi}$ | NA | 0.094 | 0.045 | 0.046 | 0.031 | NA | 0.056 | 0.031 | 0.022 | 0.020 | NA | 0.036 | 0.025 | 0.022 | 0.015 |
| | | $F_{ST}$ | NA | 0.792 | 0.407 | 0.030 | -0.002 | NA | 0.763 | 0.399 | 0.097 | 0.000 | NA | 0.732 | 0.378 | 0.101 | 0.003 |
| | 30 | $\tilde{\pi}$ | NA | 0.089 | 0.044 | 0.031 | 0.029 | NA | 0.052 | 0.029 | 0.021 | 0.018 | NA | 0.033 | 0.024 | 0.017 | 0.014 |
| | | $F_{ST}$ | NA | 0.787 | 0.421 | 0.083 | -0.002 | NA | 0.767 | 0.398 | 0.098 | 0.001 | NA | 0.750 | 0.391 | 0.115 | 0.003 |
| | 40 | $\tilde{\pi}$ | NA | 0.086 | 0.041 | 0.030 | 0.027 | NA | 0.049 | 0.029 | 0.020 | 0.018 | NA | 0.031 | 0.023 | 0.017 | 0.014 |
| | | $F_{ST}$ | NA | 0.788 | 0.411 | 0.087 | -0.003 | NA | 0.761 | 0.403 | 0.107 | 0.001 | NA | 0.720 | 0.393 | 0.115 | 0.009 |
| 5,000 | 10 | $\tilde{\pi}$ | 0.264 | 0.083 | 0.049 | 0.039 | 0.037 | NA | 0.054 | 0.033 | 0.025 | 0.023 | NA | 0.041 | 0.026 | 0.020 | 0.018 |
| | | $F_{ST}$ | 0.674 | 0.653 | 0.290 | 0.036 | -0.006 | NA | 0.637 | 0.294 | 0.079 | -0.003 | NA | 0.624 | 0.382 | 0.071 | 0.000 |
| | 20 | $\tilde{\pi}$ | NA | 0.077 | 0.046 | 0.035 | 0.033 | 0.049 | 0.049 | 0.031 | 0.023 | 0.021 | NA | 0.038 | 0.025 | 0.023 | 0.016 |
| | | $F_{ST}$ | NA | 0.645 | 0.303 | 0.042 | -0.004 | 0.638 | 0.640 | 0.302 | 0.061 | -0.005 | NA | 0.619 | 0.304 | 0.065 | 0.002 |
| | 30 | $\tilde{\pi}$ | NA | 0.071 | 0.044 | 0.033 | 0.031 | NA | 0.045 | 0.031 | 0.022 | 0.020 | NA | 0.035 | 0.024 | 0.018 | 0.015 |
| | | $F_{ST}$ | NA | 0.584 | 0.303 | 0.042 | -0.006 | NA | 0.625 | 0.30 | 0.069 | 0.001 | NA | 0.613 | 0.305 | 0.087 | 0.004 |
| | 40 | $\tilde{\pi}$ | NA | 0.072 | 0.043 | 0.031 | 0.030 | NA | 0.045 | 0.029 | 0.021 | 0.019 | NA | 0.034 | 0.023 | 0.017 | 0.015 |
| | | $F_{ST}$ | NA | 0.658 | 0.308 | 0.046 | -0.002 | NA | 0.630 | 0.306 | 0.071 | 0.005 | NA | 0.630 | 0.311 | 0.089 | 0.006 |
| 10,000 | 10 | $\tilde{\pi}$ | 0.266 | 0.079 | 0.052 | 0.040 | 0.040 | NA | 0.054 | 0.034 | 0.026 | 0.024 | NA | 0.041 | 0.026 | 0.020 | 0.018 |
| | | $F_{ST}$ | 0.665 | 0.549 | 0.213 | 0.026 | -0.004 | NA | 0.545 | 0.233 | 0.039 | -0.003 | NA | 0.529 | 0.291 | 0.052 | 0.002 |
| | 20 | $\tilde{\pi}$ | 0.261 | 0.073 | 0.045 | 0.035 | 0.035 | 0.048 | 0.048 | 0.031 | 0.023 | 0.021 | NA | 0.035 | 0.025 | 0.023 | 0.07 |
| | | $F_{ST}$ | 0.692 | 0.563 | 0.217 | 0.026 | -0.004 | 0.531 | 0.528 | 0.247 | 0.049 | -0.006 | NA | 0.522 | 0.257 | 0.041 | 0.003 |
| | 30 | $\tilde{\pi}$ | NA | 0.067 | 0.045 | 0.034 | 0.033 | NA | 0.044 | 0.030 | 0.022 | 0.021 | NA | 0.035 | 0.024 | 0.018 | 0.016 |
| | | $F_{ST}$ | NA | 0.539 | 0.233 | 0.026 | -0.005 | NA | 0.530 | 0.253 | 0.048 | -0.002 | NA | 0.524 | 0.265 | 0.066 | 0.001 |
| | 40 | $\tilde{\pi}$ | NA | 0.067 | 0.043 | 0.032 | 0.031 | NA | 0.043 | 0.030 | 0.021 | 0.020 | NA | 0.032 | 0.023 | 0.018 | 0.016 |
| | | $F_{ST}$ | NA | 0.553 | 0.242 | 0.029 | -0.008 | NA | 0.534 | 0.259 | 0.052 | -0.001 | NA | 0.525 | 0.260 | 0.073 | 0.001 |

**Figure S1.**
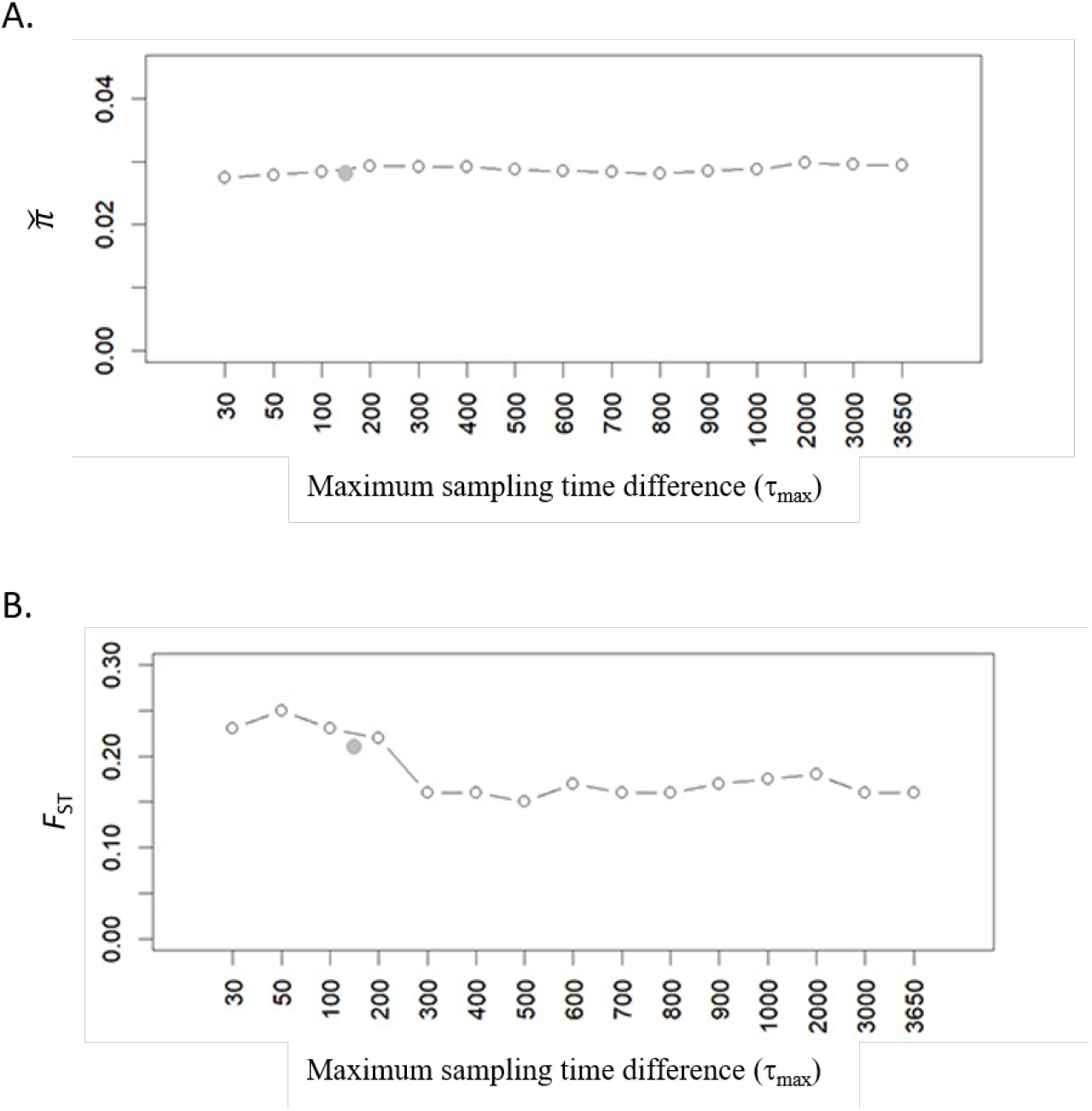
Sampling-time-corrected sequence diversity 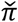 (A) and *F*_ST_ (B) of segment HA in the 10-year H3N2 data set, with the increasing upper bound of sampling time difference (τ_max_, in days) between two sequences that are compared. The gray dot indicates the result when each pair of sequences were sampled within the same 6-month flu season.

**Figure S2.**
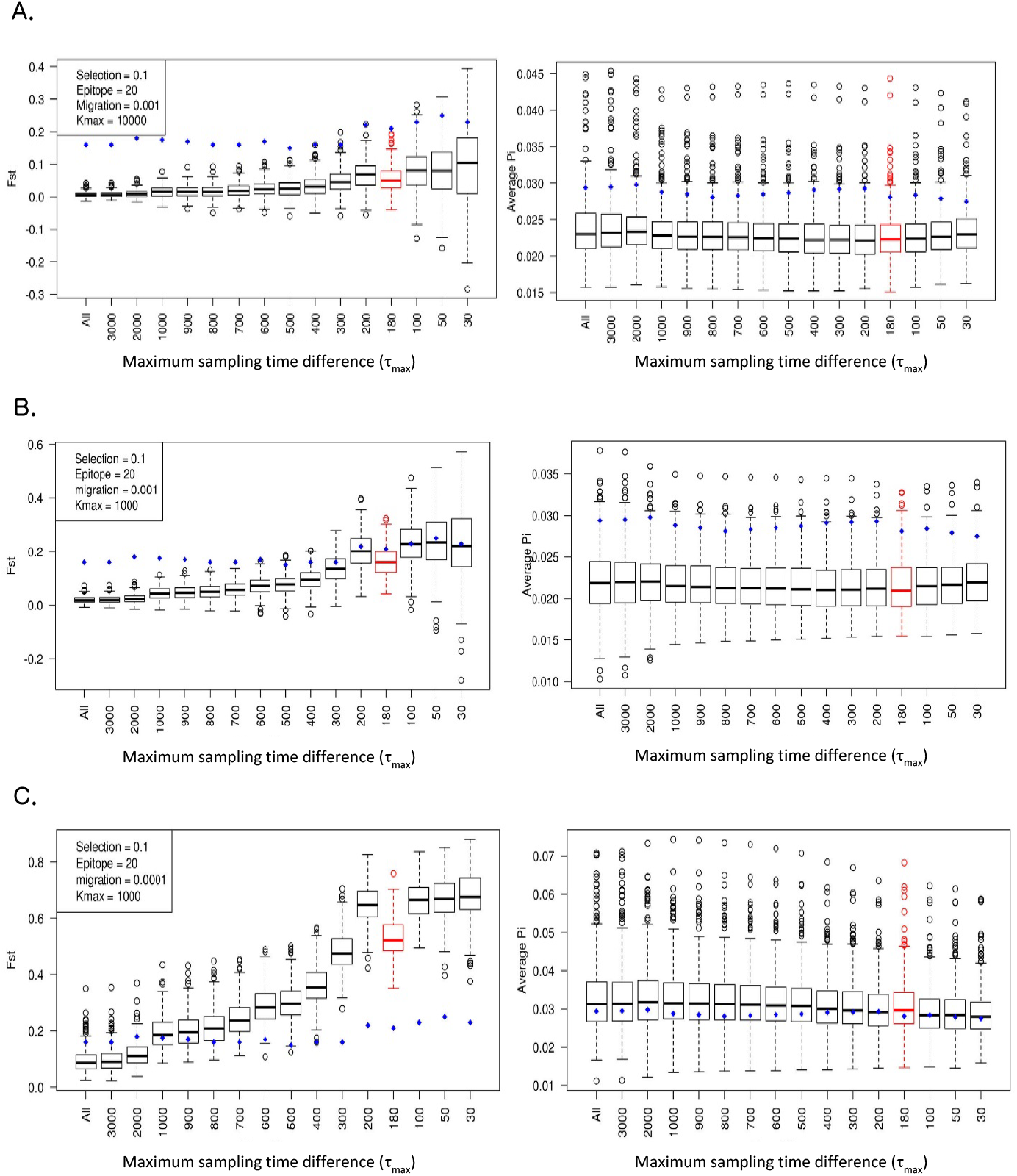
*F*_ST_ and 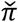 from 300 runs of simulation under positive selection (*s* = 0.1) for various τ_max_, with ε = 20 and *m* = 0.001 (A and B) or 0.0001 (C) and *K*_max_ = 1000 (B and C) or 10000 (A). The red plot indicates the result when each pair of sequences were sampled within the same 6-month flu season. The blue dots indicate the values observed in H3N2 data.

**Figure S3:**
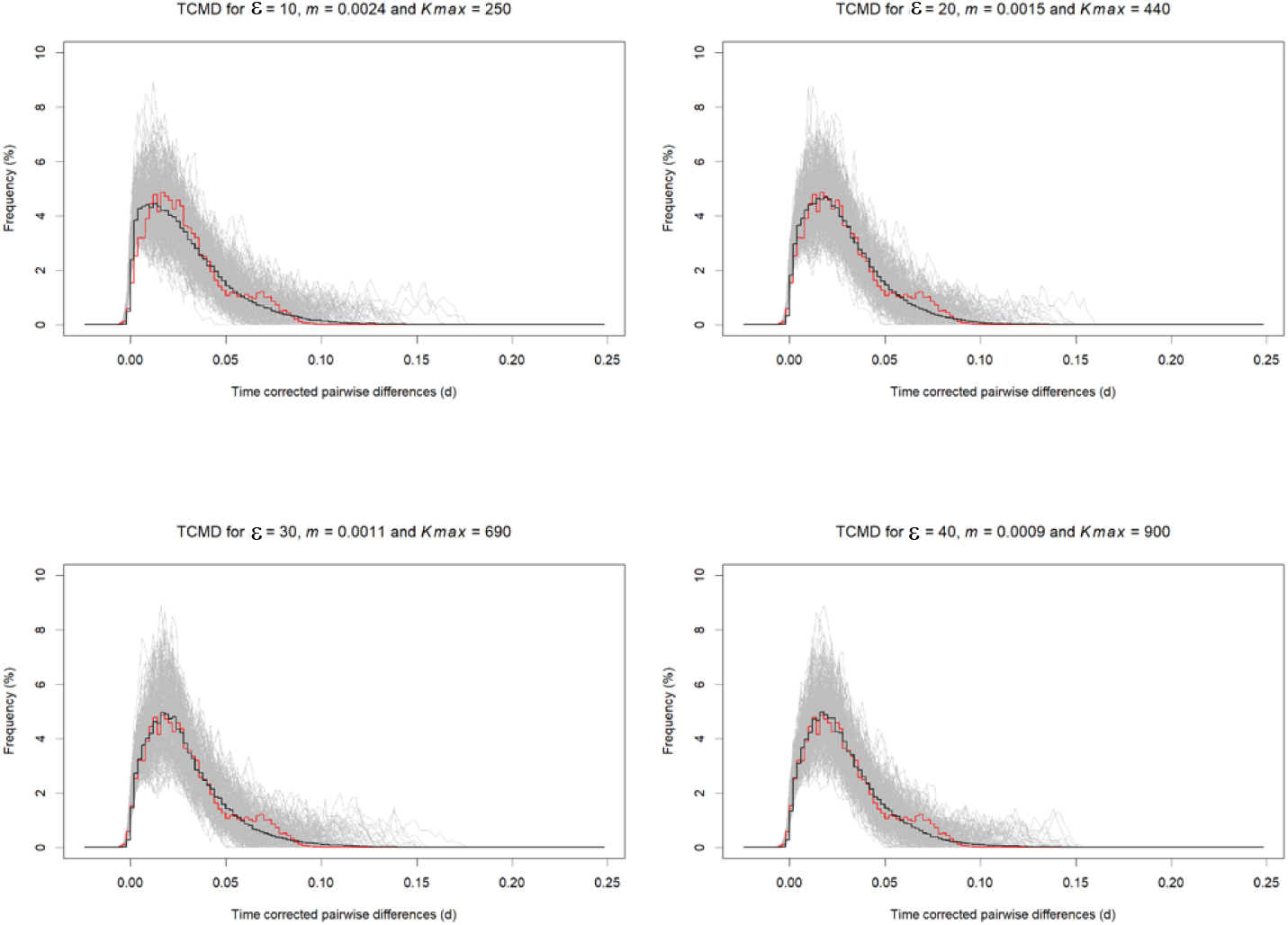
TCMD for simulated data under positive selection with *s* = 0.05 and ε = 10, 20, 30, and 40. For each set of *s* and *ε*, the best values of *K*_max_ and *m* were found as shown in Table 2. The grey curves represent 300 simulated replicates for the same parameters. The black curve represents the average of those replicates and the red one represents the observed data.

**Figure S4.**
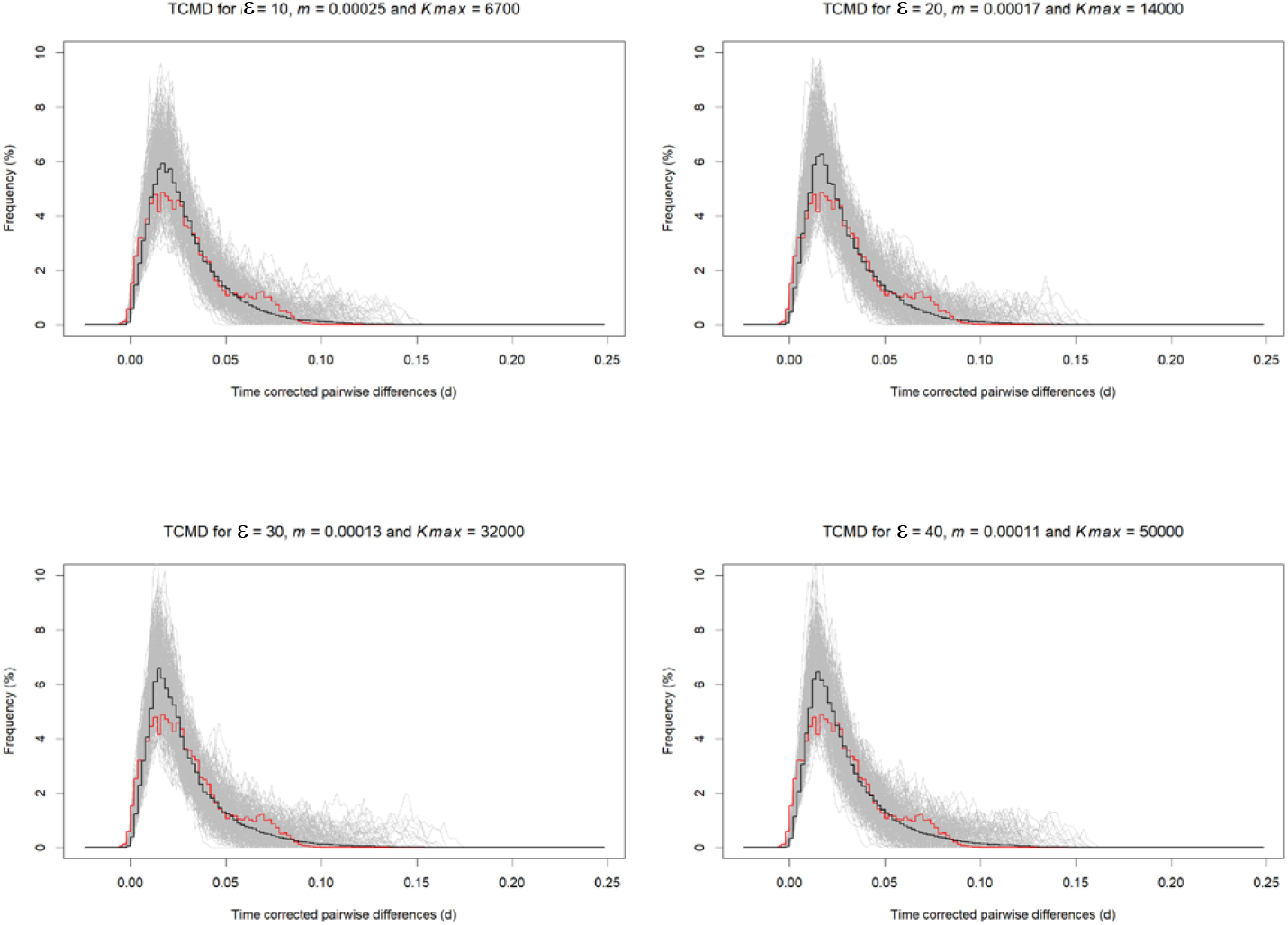
TCMD for simulated data under positive selection with *s* = 0.1 and ε = 10, 20, 30, and 40. For each set of *s* and ε, the best values of *K*_max_ and *m* were found as shown in Table 2. The grey curves represent 300 simulated replicates for the same parameters. The black curve represents the average of those replicates and the red one represents the observed data.

**Figure S5.**
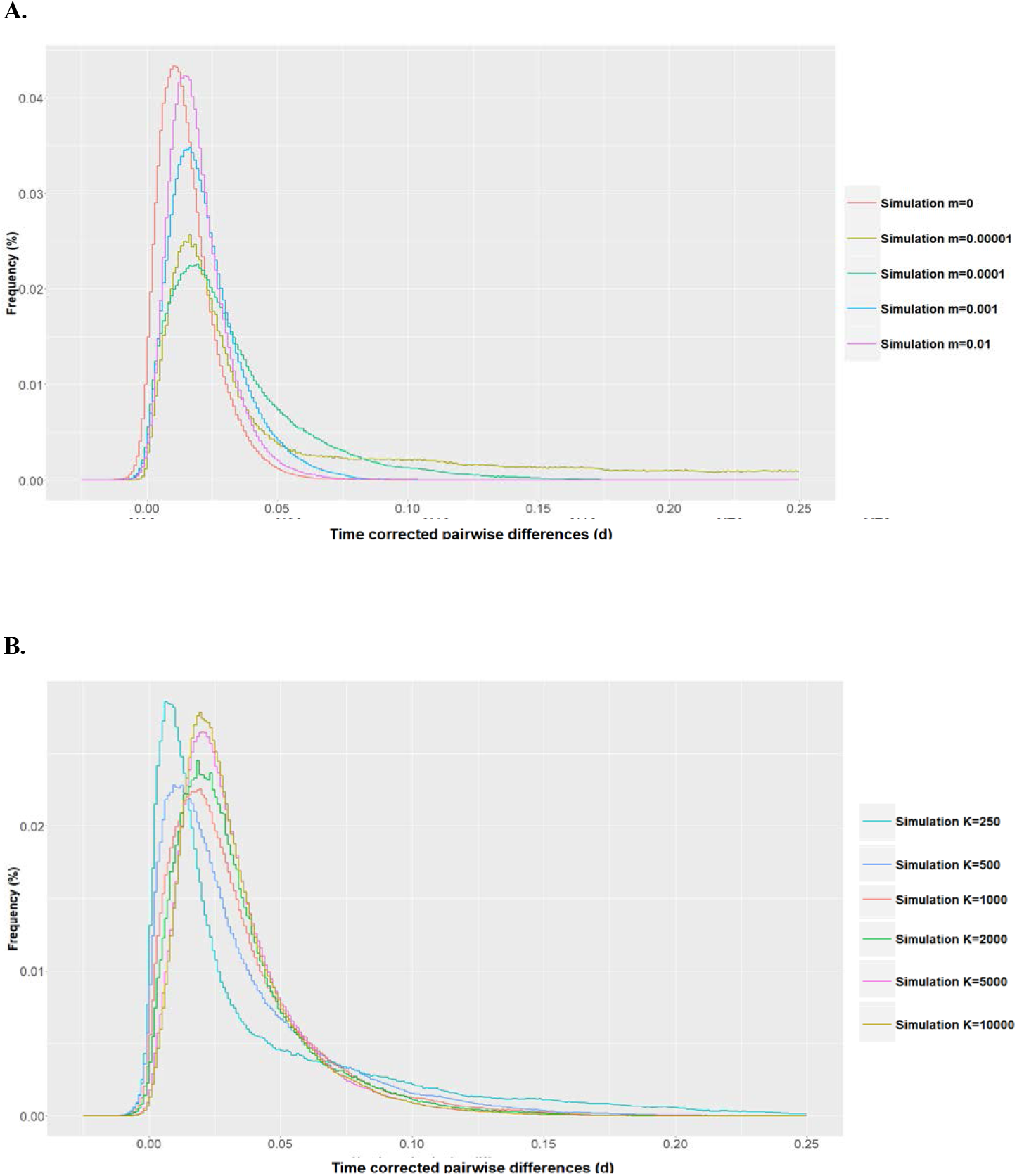
TCMD of simulated 10-year HA sequences for different values of *m* (**A**; with *K*_max_ = 1000) and of *K*_max_ (**B**; with *m* = 0.0001). Other parameters: *s* = 0.1 and ε = 10. Each curve is an average over 300 simulation runs.

**Figure S6.**
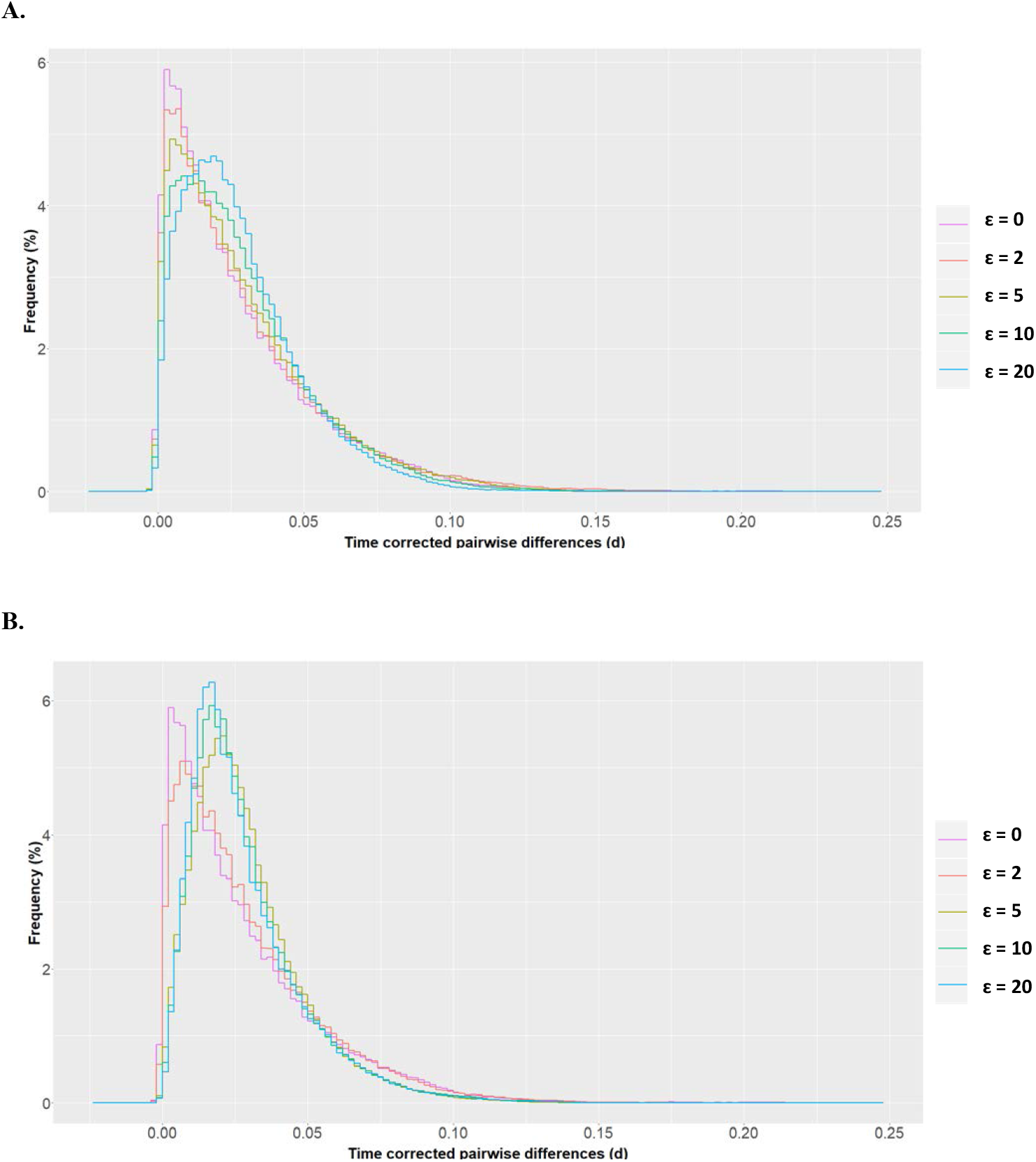
TCMD for simulated data with increasing frequency of beneficial mutations (ε) and other parameters that yield π and *F*_ST_ matching the observed data. Under neutrality (ε = 0), *m* = 0.004 and *K*_max_ = 110. Under positive selection with *s* = 0.05 (**A**), we use (ε, *m*, *K*_max_) = (2, 0.0036, 130), (5, 0.003, 170), (10, 0.0024, 250), and (20, 0.0015, 440). With *s* = 0.1 (**B**), we use (ε, *m*, *K*_max_) = (2, 0.003, 180), (5, 0.0004, 2500), (10, 0.00025, 6700), and (20, 0.00017, 1400).

**Figure S7.**
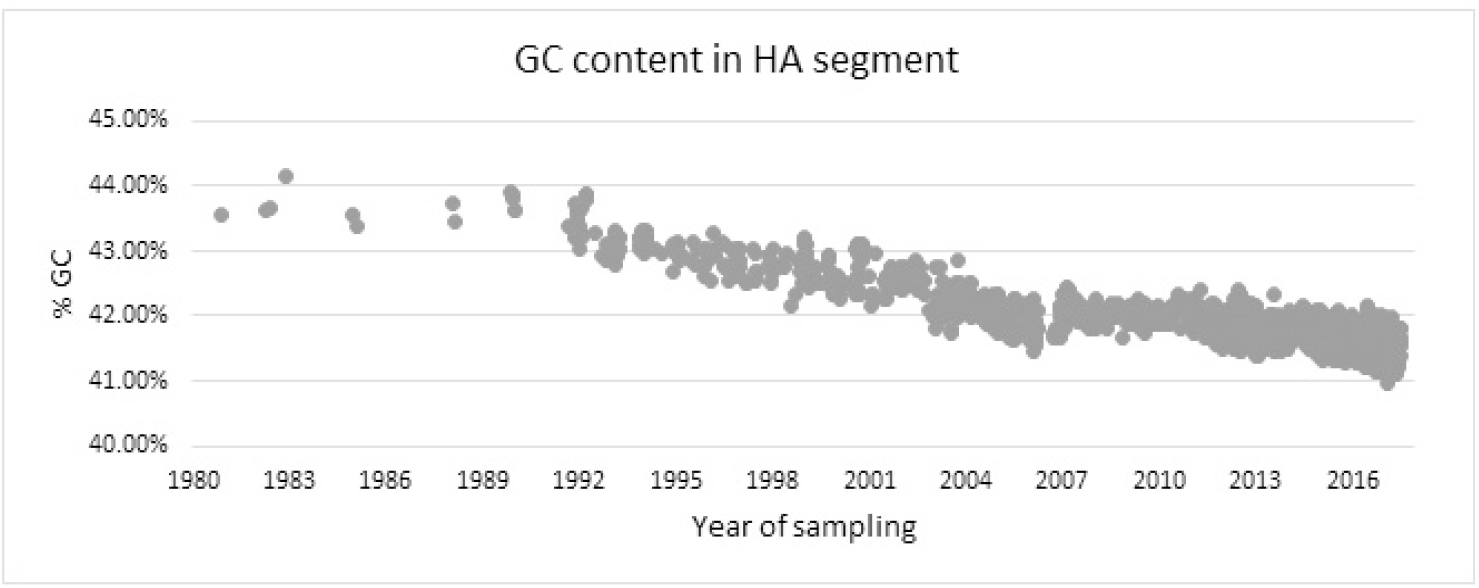
Plot of the GC content (in %) of the HA segment of H3N2 viruses against the time of sampling.

